# Design nonrepetitive and diverse activity single-guide RNA by deep learning

**DOI:** 10.1101/2024.05.30.596019

**Authors:** Yan Xia, Zeyu Liang, Xiaowen Du, Dengtian Cao, Jing Li, Lichao Sun, Yi-Xin Huo, Shuyuan Guo

## Abstract

Multiplex and precise control of the gene expression based on CRISPR/Cas9 is important to metabolic regulation in synthetic biology. However, employing single guide RNAs (sgRNAs) that possess repetitive DNA sequences and exhibit uniform activity could detrimentally affect the editing process, undermining both its stability and regulatory potential. In this study, we developed a deep generative model based on a decoder-only Transformer architecture (sgRNAGen) for the *de novo* generation of a series of nonrepetitive and diverse sgRNAs with activity. To assess the quality of sgRNAs generated by sgRNAGen, we evaluated their activity by targeting essential genes, with the results indicating that 98% of the generated sgRNAs were active in *Bacillus subtilis*. The generated sgRNAs were further validated for applications in single-gene editing, large fragment knockouts, and multiplex editing. Notably, the efficiency of knocking out long fragments up to 169.5 kb reached 100%, and targeting multiple sites allowed for the creation of strains with various combinations of mutations in a single editing. Furthermore, we developed a CRISPRi system utilizing the designed sgRNAs to regulate gene expression with desired strength and high precision. SgRNAGen offers a method for devising nonrepetitive and diverse activity sgRNAs, enhancing metabolic control and advancing applications within synthetic biology.

**TOC:** 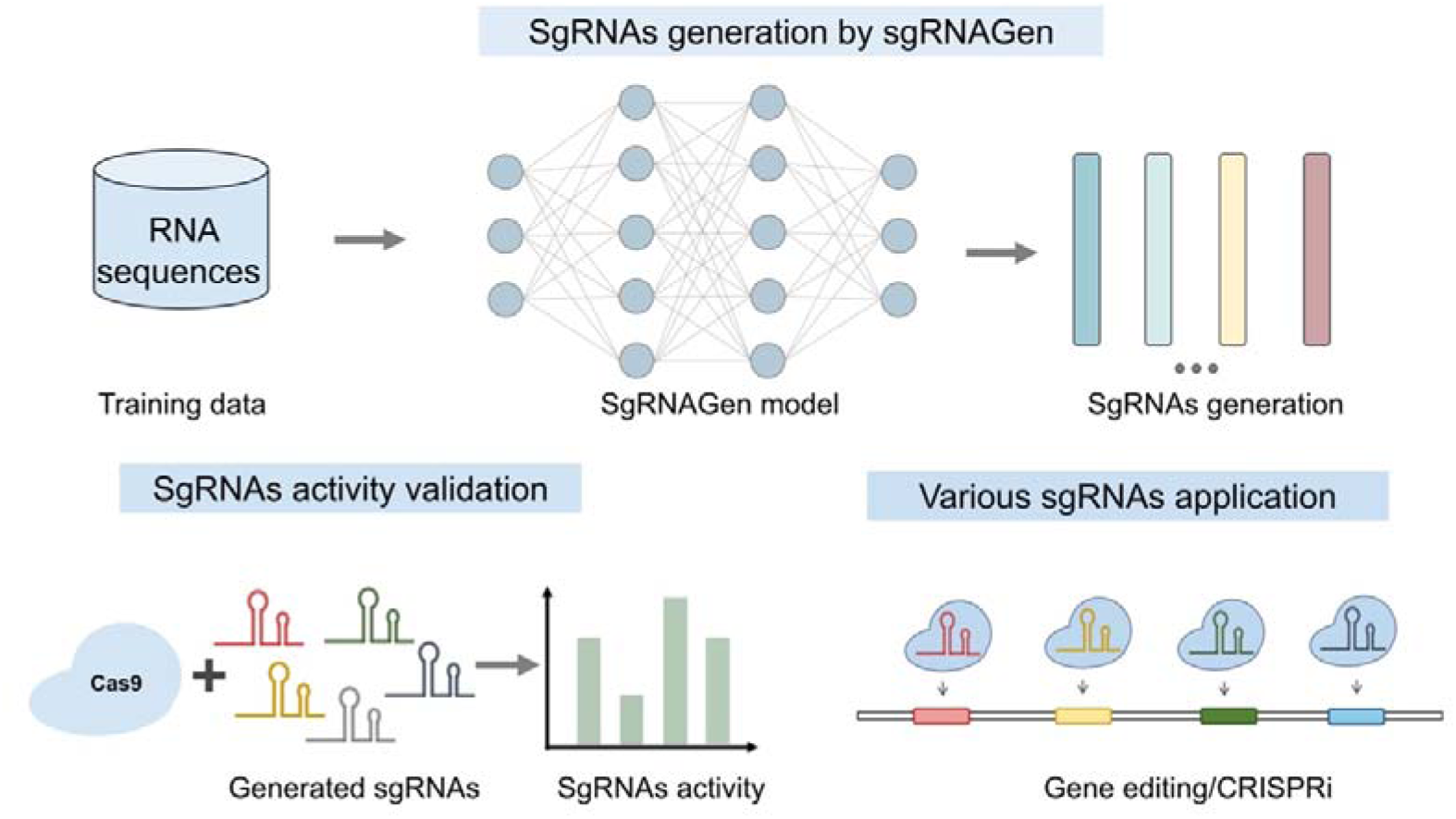

## INTRODUCTION

The Type II Clustered Regularly Interspaced Short Palindromic Repeats (CRISPR)/Cas9 system has emerged as a highly efficient gene editing tool^1–4^. Cas9 from *Streptococcus pyogenes* (SpCas9) comprises the Cas9 protein, CRISPR RNA (crRNA), and trans-activating CRISPR RNA (tracrRNA)^5, 6^. The crRNA, containing a 20-nucleotide spacer sequence, collaborates with tracrRNA and the Cas9 protein to form a ternary complex, which targets complementary DNA sequences adjacent to a 3-base pair sequence known as the NGG protospacer adjacent motif (PAM)^7, 8^. Upon binding, it induces cleavage of the double-stranded DNA, resulting in a specific break. To simplify this system, crRNA, and tracrRNA have been merged into a single, chimeric single-guide RNA (sgRNA) using a GAAA linker^7^. This sgRNA guides the Cas9 enzyme to perform sequence-specific cleavage of double-stranded DNA with high efficiency.

Due to the complexity of intracellular metabolic networks, achieving precise control over metabolic flux through multiplex gene editing is crucial for the construction of robust microbial chassis^9–11^. The CRISPR/Cas9 has been developed the powerful tools in multiplex genome editing and transcriptional regulation^9^. For example, utilizing the CRISPR/Cas9-facilitated Multiplex Pathway Optimization (CFPO) technique, a strain exhibiting a three-fold enhancement in xylose-utilization rate was developed, achieved through the simultaneous regulation of four genes^12^. The technology of multiplex base editing was employed to modulate the metabolic flux of acetoin production in *Bacillus subtilis*, establishing a library of gene modifications in competitive and central metabolic pathways, successfully enhancing the yield by 26.3%^13^. CRISPR interference (CRISPRi) enables precise fine-tuning of repression levels, a method instrumental in balancing the metabolic network and enhancing production^14, 15^. By employing multiplex combinatorial knockdown of competing pathways through CRISPRi, the isopentenyl titer was successfully increased by up to 98% in *Escherichia coli*^16^. Targeting multiplex sites often need to co-expressed multiple sgRNAs, which contained several long DNA repeats in sgRNAs expression box. However, DNA sequences with repeats presents significant challenges *in vitro* assembly, including mutations and deletions, and leads to genetic instability *in vivo*, especially in organisms with robust homologous recombination systems^17–19^. In addition, it is still hard to precisely repress the gene expression level by a specific sgRNA^20^.

Some studies aimed to create non-repetitive and various activities sgRNAs by random mutagenesis or iterative trial-and-error. Byun *et al.* developed an sgRNA library through random mutagenesis, specifically targeting the tetraloop and its adjacent regions, yielding a collection of mutant sgRNAs^20^. Remarkably, these sgRNAs demonstrated repression efficiencies at the transcriptional level that were up to 45 times greater compared to their non-repressing counterparts. However, the similarity between variants and wild type sgRNA are high due to the mutants only appear in certain areas. Additionally, a rational design approach was employed to generate non-repetitive sgRNAs^21^. The method of Extra-Long sgRNA Arrays (ELSAs) was developed, enabling the simultaneous and stable expression of 28 sgRNAs. This was achieved by analyzing the biochemical characteristics and biophysical modeling of the sgRNAs. ELSAs were then utilized to co-express 22 sgRNAs, resulting in the repression of 13 genes by up to 3500-fold. However, designing sgRNAs through ELSAs requires multiple rounds of experimental validation to characterize more nonrepetitive sgRNA.

Given the vastness and complexity of RNA sequence space, which increases exponentially with sequence length, the experimental search for specific RNA sequences remains costly and inefficient^22, 23^. Therefore, there is an urgent need for a comprehensive computational platform to thoroughly explore the sequence space and efficiently design functional RNAs. With the advancement of deep neural networks in computational biology, they have demonstrated immense potential in prediction and *de novo* RNA generation tasks, such as riboswitches, terminators, tRNAs, ribozymes and others^22, 24–26^. For example, *Sumi et al.* designed the RNA Family Sequence Generator (RfamGen), a novel approach for generating functional RNA family sequences, which involves sampling points from a semantically rich and continuous representation, enabling the design of artificial sequences^27^. Remarkably, these artificially designed sequences exhibit higher activity than their natural counterparts, showing deep generative models have exhibited the great potential in RNA design^28, 29^.

In this study, to design novel and non-repetitive sgRNAs, we designed and trained a deep learning generative model based on the Transformer architecture (sgRNAGen). These generated sgRNAs were crafted with strict constraints on ΔG and RNA structure to achieve high activity levels. We generated sgRNAs by sgRNAGen and tested fifty sgRNAs, targeting essential genes in *B. subtilis*, to evaluate their binding affinity with the Cas9 protein. Furthermore, to verify sgRNAs gene-editing efficiency, we randomly selected eight sgRNAs for the targeted knockout of the non-essential *bdhA* gene, with editing efficiencies ranging from 12.5% to 75%. In addition, the sgRNAs demonstrated the capability to knockout large genomic fragments, varying from 21.3 kb to 169.5 kb, with efficiencies ranging from 12.5% to 100%. We extended our investigation to multiplex gene editing, using the designed sgRNAs to target two or three sites simultaneously. This approach successfully produced various combination strains through a single editing event. The designed sgRNAs were also employed in the CRISPRi system for the fine-tuning of gene expression levels. By randomly selecting 14 sgRNAs targeting sfGFP, we achieved suppression of sfGFP expression levels within the range of 34.6% to 73.4%, which can be utilized to accelerate the optimization of metabolic pathways. Our model could be effectively used to design novel and non-repetitive sgRNAs for metabolic regulation and synthetic biology applications.

## RESULTS

### A generative model based on Transformer trained to generate sgRNAs

Deep learning has been extensively applied to CRISPR-related applications, including off-target prediction, sgRNA activity, and editing outcome prediction, typically relying on supervised learning models^30^. However, the lack of sufficient data presents significant challenges for designing new crRNAs using these models. To address the challenge of limited experimental data, we employed an unsupervised pre-training approach. This approach has the advantage of learning the underlying distribution of the samples without requiring annotation^31, 32^. Once the true sample distribution function is learned, samples can be generated from scratch using autoregressive methods or optimized using techniques like Gibbs sampling. The Transformer model is good at this because it can understand long-distance relationships in text using its self-attention mechanism. Building on this strength, we have developed a decoder-only Transformer-based model (sgRNAGen) specifically designed to discern the unique characteristics of CRISPR-related sgRNA sequences.

In the CRISPRCas-db metagenomics database, over 40,000 CRISPR-related RNA sequences have been utilized for training the sgRNAGen model. RNA sequences are tokenized at the character level in sgRNAGen, maintaining the original positioning of nucleotides in the sequence and thus enhancing the control over RNA generation (**Figure 1C**). The sgRNAGen was trained using an autoregressive approach, where the probability of the occurrence of the next base in the sequence solely depends on the preceding nucleotide. The performance of the sgRNAGen model was evaluated and refined through the utilization of cross-entropy loss, which quantifies the discrepancy between the probability distribution of the model predictions and the probability distribution of the true labels. The sgRNAGen model updates its numerical representation through GPT2Blocks, employing multi-head self-attention mechanisms (**Figure 1D**). Generative models are often plagued by hallucinations, where the generated samples conform to the model’s fitted distribution but are not realistic. Therefore, additional methods are needed to exclude these unrealistic samples. Given the critical importance of secondary structure stability for the functionality of RNA molecules, we incorporate secondary structure prediction and energy calculation to constrain the samples designed by the model. Additionally, to address the divergence issue encountered by autoregressive models during generation where an error in predicting a single token can lead to increasing errors in subsequent tokens, we employ a position-agnostic sampling strategy. Specifically, we randomly select a mutation in the initial sequence and use the trained GPT model to score it, accepting sequences that show a decrease in loss. The ΔG value was used as a criterion for evaluation during sequence generation: mutations are accepted when the ΔG exceeds a specific threshold. Otherwise, the sequence is regenerated. Additionally, considering the close correlation between the activity of sgRNA and its secondary structure, we ensure that the generated sgRNA sequences have secondary structures similar to that of the wild-type (**Figure 1E**). 10% of the dataset sequences were reserved for evaluation and were not included in the training process.

**Figure 1.**
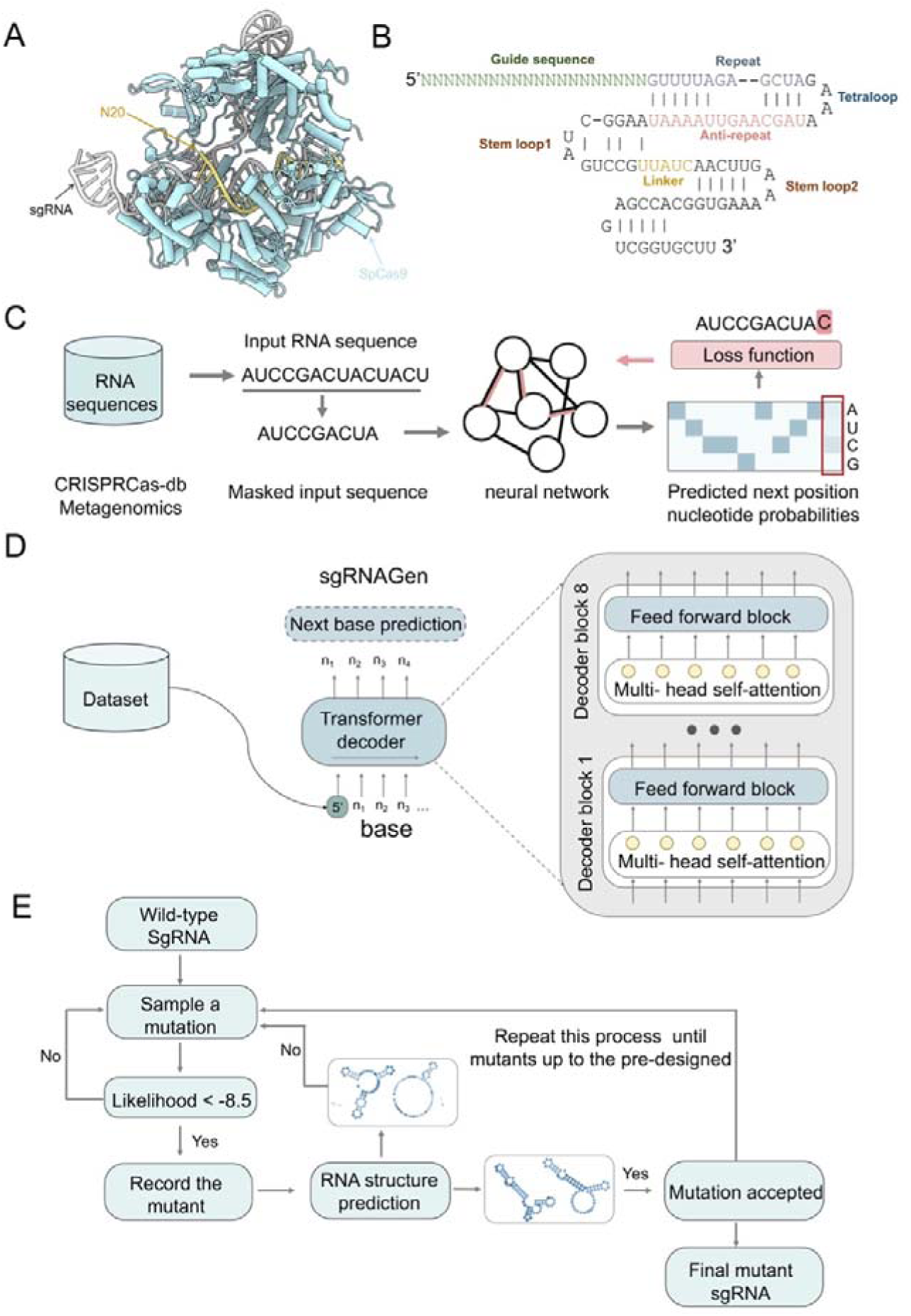
A framework for design the sgRNAs by deep learning. (A) Formation of a ternary complex by the sgRNA, N20 sequence, and SpCas9 protein. (B) Schematic representation of the sgRNA and N20 sequence. (C-D) Illustration of the generative model architecture for sgRNAs. (E) Workflow diagram for the design of sgRNAs.

### The activity validation of generated sgRNAs in *B. subtilis*

To validate the effectiveness of sgRNAGen model, we *de novo* generated 50 sgRNA sequences of 61 nt in length using our trained model. These sequences featured various numbers of mutated bases, specifically 8, 10, 12, 14, and 24. Additionally, previous research has indicated that positions 7, 23, 24, 31, and 33 in sgRNA are critical sites, where mutations can significantly reduce activity. Therefore, we maintained the nucleotides at these five positions unchanged during the sequence generation process. Subsequently, experimental validation was conducted in *B. subtilis* for these synthesized sequences. We employed a simple growth-based selection system to assess the activity of sgRNA in *B. subtilis*. The experimental design targeted the essential gene *secY*, evaluating activity by replacing different sgRNA sequences. If the sgRNA is active, it will guide Cas9 to cut the essential gene in the host cell, resulting in the absence of colonies on the plate. Conversely, if the sgRNA is inactive, colonies will grow on the plate (**Figure 2A**). Wild-type sgRNA and a no-sgRNA condition were used as positive and negative controls, respectively.

**Figure 2.**
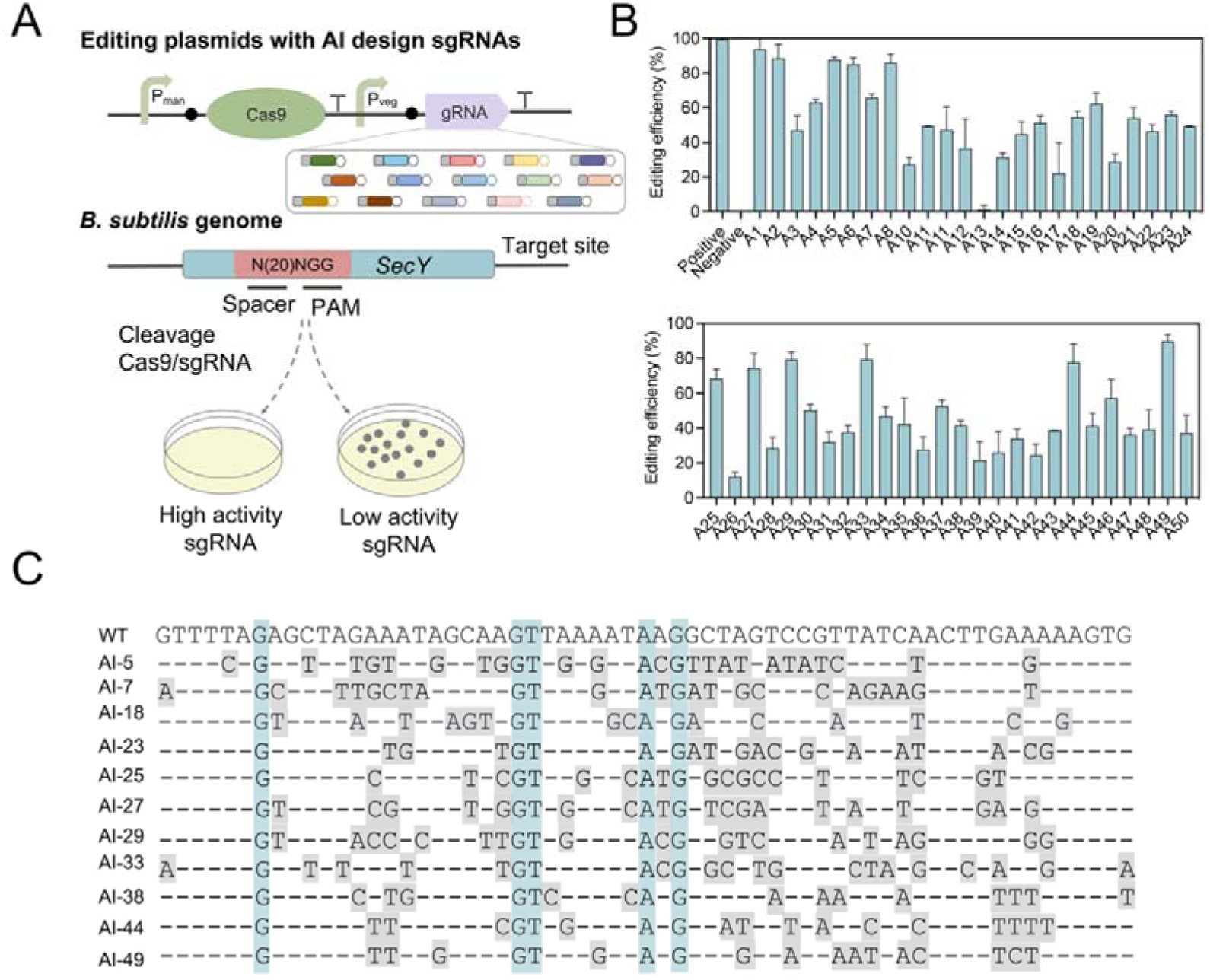
Assessment of the generated sgRNAs activities in *B. subtilis*. (A) Schematic representation of the methodology for assessing sgRNA targeting efficiency using a growth-based selection assay. (B) Relative cleavage activity of the generated sgRNAs (n = 3 independent replicates). (C) Segments of the generated sgRNA sequences. Horizontal lines denote sequences identical to the wild-type sgRNA. The five bases highlighted against a blue background are fixed and remain unaltered, while bases on a gray background represent mutations.

As shown in figure 2B, nearly all the generated sgRNAs exhibit activities, with the exception of A13. Notably, eight sgRNAs have at least 80% of the activity compared to wild-type sgRNAs, specifically A1, A2, A5, A6, A8, A33, A44, and A49. Additionally, approximately half of the generated sgRNAs retain at least 50% activity. These results demonstrate that our model has successfully learned the conformation and distribution relevant to CRISPR-associated RNA, enabling it to generate active sgRNAs. Upon analyzing the activities and the number of sgRNA mutants, we observed no correlation between them (**Figure 2B and 2C**). For instance, A13, which has 11 mutants, exhibits almost no activity, whereas A5 and A29, with 22 and 18 mutants respectively, maintain 87.7% and 74.7% activity. It is as expected that the impact of each mutation of sgRNA is different and need to consider the whole structure changes. Furthermore, Additionally, we employed the online RNA structure prediction platform, RNAfold, to ascertain the secondary structure of the generated sgRNAs. The secondary structures of these sgRNAs closely resemble those of the wild-type sgRNA, aligning with our design principles (**Supplementary Figure S1**).

### Gene editing by generated sgRNAs

CRISPR/Cas9 has been widely used in genome editing, and we evaluated the gene editing efficiency of generated sgRNAs by deep learning. The non-essential gene *bdhA* was used as the targeting sites. Eight sgRNAs - A3, A4, A7, A15, A37, A38, A44, and A49 - were arbitrarily selected for editing the target site with the same N20 sequence, with the wild-type sgRNA serving as a positive control (**Figure 3A**). We assessed the editing efficiency through colony PCR and Sanger sequencing, selecting eight colonies from each plate for analysis. The results showed that all designed sgRNAs effectively knocked out the target genes, confirming their potential for gene editing in future experiments. However, the editing efficiencies varied among the different sgRNAs. For example, A49 achieved the highest editing efficiency at 75%, while A3 and A38 had the lowest, each at 12.5% (**Figure 3B**).

**Figure 3.**
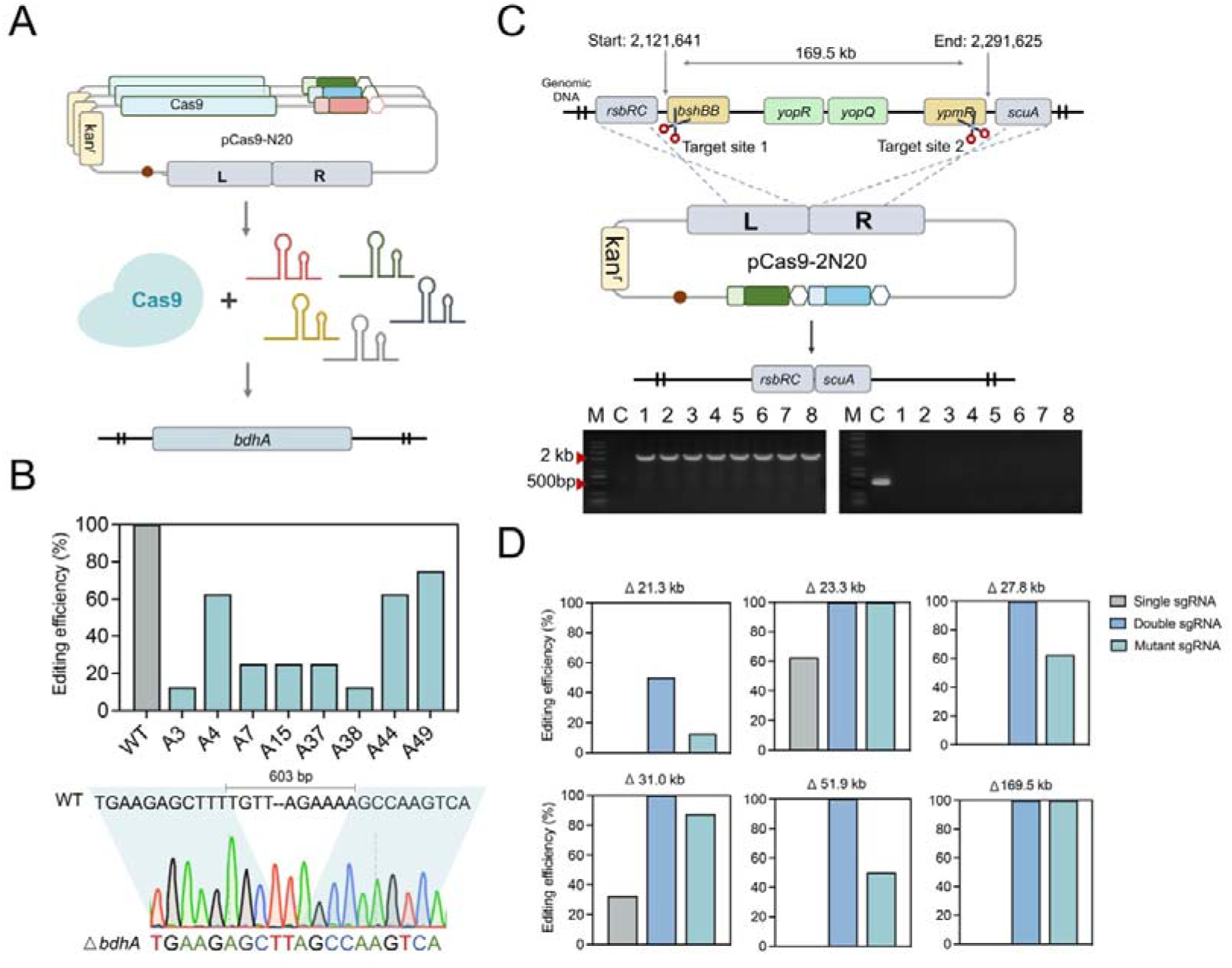
The generated sgRNAs mediated genome editing in *B. subtilis*. (A) Schematic for the targeting of the *bdhA* gene using various sgRNAs. (B) Editing efficiency of the generated sgRNAs, with positive colonies selected for Sanger sequencing. (C) Schematic representation of large-scale fragments knockout, where two N20 sequences were designed for target cleavage. (D) Editing efficiency of large-scale knockouts ranging from 21.3 kb to 169.5 kb.

To optimize the efficiency of microbial cell factories, strategies often involve the deletion of redundant, non-essential regions to reduce transcription costs and eliminate competing pathways. Rapid knockout of these non-essential regions relies on the effectiveness of technologies capable of long fragment deletions. Our previous research demonstrated that expressing two sgRNAs enhances the capacity for large fragment knockout via CRISPR/Cas9 in *B. subtilis*. Nevertheless, constructing plasmids for this knockout system is time-consuming and labor-intensive, primarily due to the repetitive sequences of sgRNAs. To circumvent this issue of repetitive sequences, we employed diverse sgRNAs with non-repetitive sequences for editing large fragments, thereby avoiding DNA repeat-induced plasmid instability and assembly challenges. Each of the two sgRNAs was specifically designed to target genes situated at either end of the large fragment for knockout (**Figure 3C**). We evaluated the effectiveness of the generated sgRNAs on six large-fragments, with the sizes of the knockout segments being 21.3 kb, 23.3 kb, 27.8 kb, 31.0 kb, 51.9 kb, and 169.5 kb, respectively. In addition, we used the knockout efficiency of both single and dual wild-type sgRNAs as controls. The former struggled with large fragment editing, consistent with our prior findings, while the latter achieved high efficiency in large fragment knockout. The results of diverse sgRNAs indicated that all targeted large fragments were successfully knocked out with efficiencies ranging from 12.5% to 100% (**Figure 3D**). For example, the efficiency of 169.5 kb large fragments knockout was 0%, 100%, and 100% using single wild-type sgRNA, dual wild-type sgRNAs, and designed sgRNAs, respectively. This validated demonstrates that the sgRNAs we generated are effective in enabling large fragment knockouts, thereby streamlining the process of plasmid construction to necessitate only a single-cycle approach.

### Multiplex gene editing using generated sgRNAs

To construct efficient microbial chassis, multiple rounds of gene editing are typically required to rewire metabolic pathways. Although the use of multiplex gene editing techniques significantly reduces the labor and time involved in manipulating multiple genes, thereby accelerating the development of the desired chassis, this process is challenges^21^. Specifically, when employing the CRISPR/Cas9 system for multiplex gene editing, the repeated use of sgRNAs complicates and destabilizes plasmid construction, thereby increasing the complexity of the procedure. Using diverse sgRNAs could easily express multiple sgRNAs due to the low homologous.

To verify the multiplex editing capacity of the generated sgRNAs, we utilized the designed sgRNAs targeting two or three sites to knockout these gene, simultaneously. Containing various sgRNAs with non-repetitive DNA sequences could be built by one round. We synthesized a series primers and assembly them *in vitro* by PCR to achieve two, three, and four sgRNA expression cassettes, respectively. The sgRNAs of A4, A44, and A48 were used to multiple gene editing. The related repair templates by PCR using *B. subtilis* genome as template with sgRNA expression cassettes to Gibson assembly (**Figure 4A**). We randomly selected the genes *nprE* and *wprA* for knockout experiments. The editing efficiencies for *nprE* and *wprA* were found to be 87.5% and 62.5%, respectively (**Figure 4B**). The combined efficiency for the knockout of both genes was up to 62.5%. Similarity, we knockout the genes *wprA* and *epr*, the simultaneous knockout efficiency was 50% (**Figure 4C**). We subsequently attempted the simultaneous knockout of two large fragments. The pairs of fragments, measuring 17.8 kb and 19.6 kb, and 19.6 kb and 23.3 kb, were targeted for knockout using designed sgRNAs. The results indicated that it was feasible to concurrently knockout both sets of large fragments, with knockout efficiencies of 32.5% and 12.5%, respectively (**Figure 4D, Supplementary Figure S2**). We further explored the effects of simultaneous knockouts of three genes including *wprA*, *epr*, and *nprE*. After gene editing, there was a significant decrease in the number of colonies on the plates, which might be due to a high cell mortality rate resulting from multiple cuts in the genome. To assess the editing efficiency at these three genomic sites, we selected 10 colonies for PCR validation. The findings revealed that the editing efficiencies for *wprA*, *epr*, and *nprE* were 70%, 30%, and 50%, respectively. Notably, 50% of the colonies exhibited editing at two genomic sites, and three combinations of two-site edits were observed. However, we did not find any strains with simultaneous editing at all three sites (**Figure 4E, Supplementary Figure S3**). This phenomenon may be due to the limited efficiency of the homologous recombination system in *B. subtilis*, leading to the death of strains with concurrent edits at three genomic sites due to ineffective repair.

**Figure 4.**
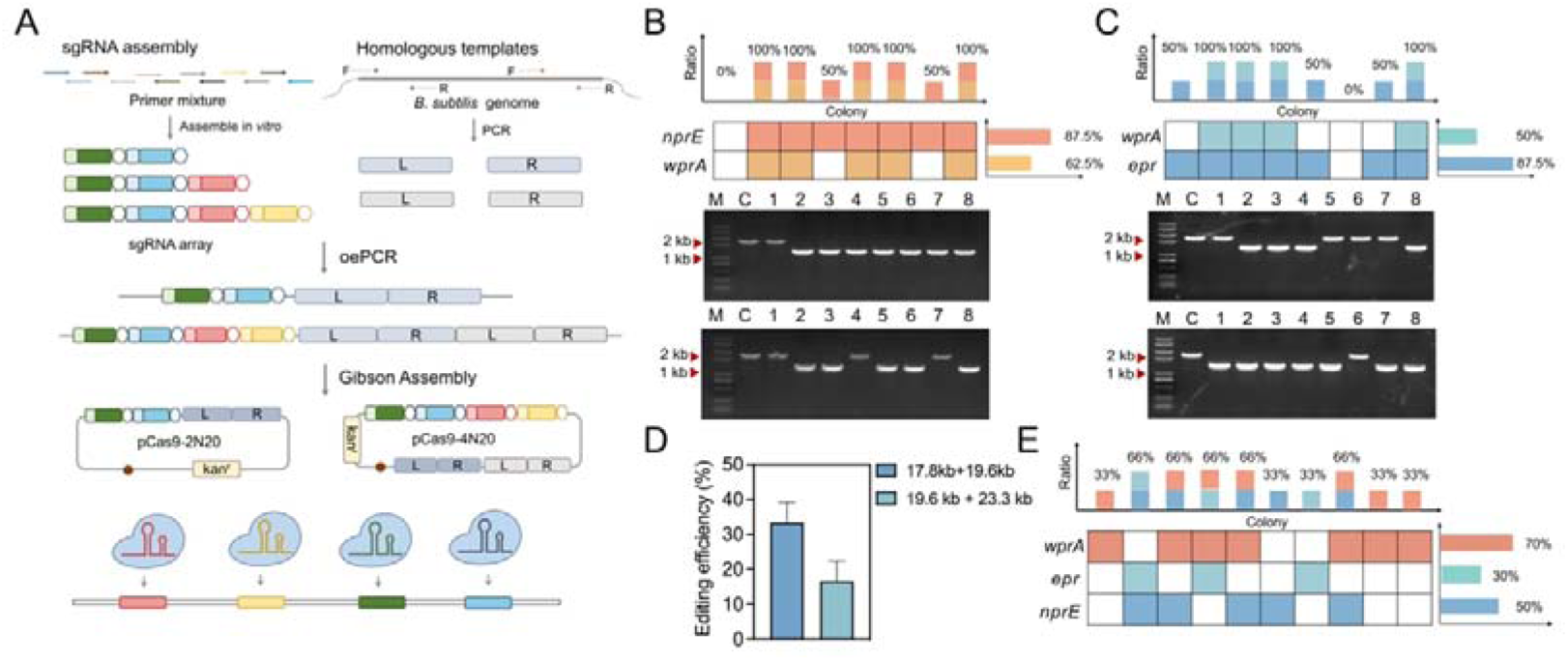
Multiplex gene editing utilizing generated sgRNAs in *B. subtilis*. (A) Schematic depicting the rapid and efficient assembly of multiple sgRNAs. (B) Editing efficiency of simultaneously knocking out *nprE* and *wprA*. (C) Editing efficiency of simultaneously knocking out *wprA* and *epr*. (D) Editing efficiency of knocking out two large-scale fragments. (e) Editing efficiency of simultaneously knocking out *wprA*, *epr*, and *nprE*.

### CRISPRi system inhibition efficiency of generated sgRNA

In metabolic engineering, the rational allocation of resources to reshape metabolic pathways is of paramount importance. Completely inhibiting the by-pathways may not always be the optimal strategy. Fine-tuning gene expression levels can help avoid the accumulation of intermediates and toxic substances, thereby achieving a balance between growth and production pathways^20^. The designed sgRNAs with varying activities can be utilized for the fine-tuning of gene expression via the CRISPRi system, employing the dead SpRY, a Cas9 variant with PAM-less specificity. To validate the repression efficiency of different sgRNAs, we employed a fluorescence reporting system based on the sfGFP gene (**Figure 5A**). The dSpRY-sgRNA-DNA ternary complex targets the transcriptional region, inhibiting RNA polymerase transcription the reporter gene *sfGFP*. The repression efficiency of the CRISPRi system is reflected in the fluorescence intensity, where lower fluorescence levels indicate higher inhibition efficiency.

**Figure 5.**
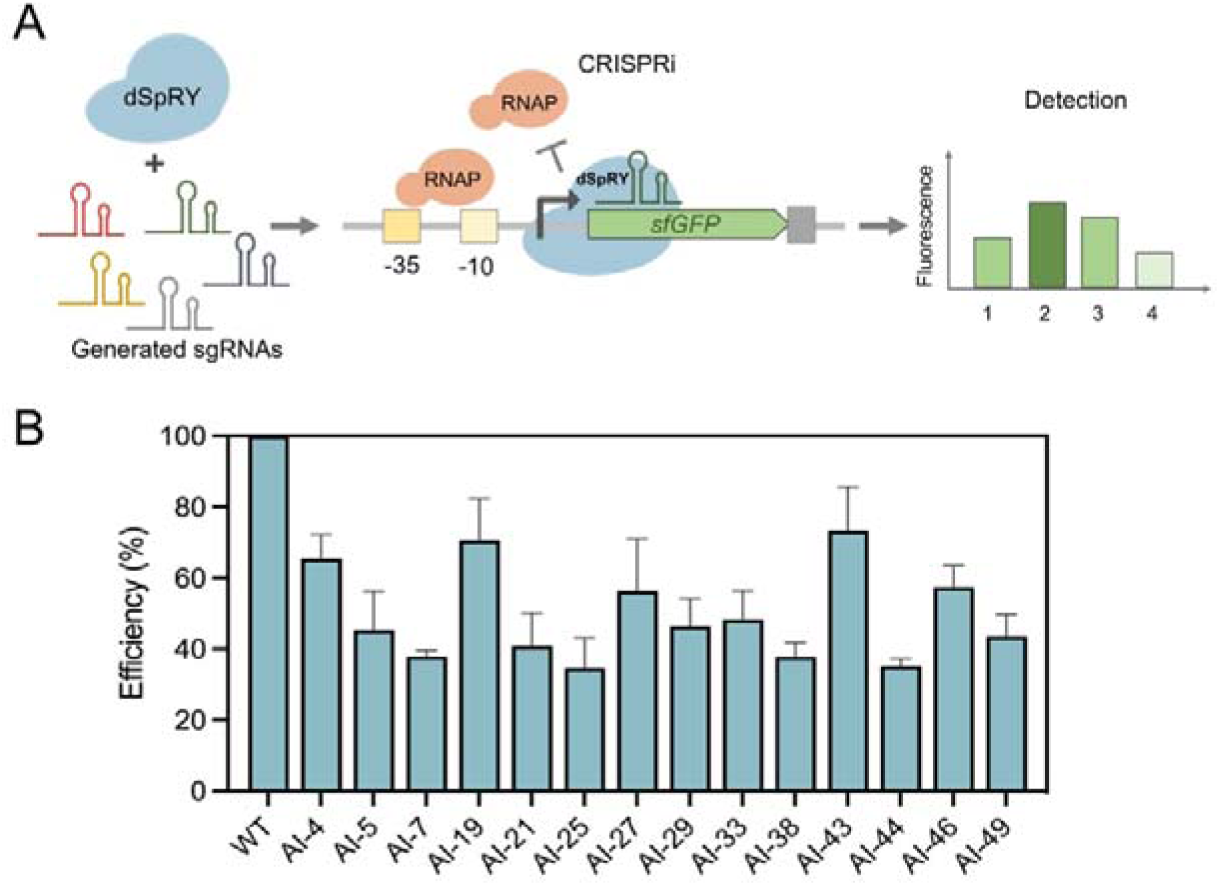
Evaluation of the repression efficiency of generated sgRNAs. (A) The detection system for assessing repression efficiency. (B) Repression efficiency of the generated sgRNAs (n = 3 independent replicates).

To identify the optimal target site for gene repression, we designed four sites within the promoter region inhibiting sfGFP expression using wild type sgRNA. Among these, target site 3 demonstrated the highest efficiency (**Supplementary Figure S4**). Consequently, we selected 14 sgRNAs targeting site 3 to evaluate their repression efficiency, using the wild type sgRNA as positive contorl. These sgRNAs exhibited a range of inhibitory activities on sfGFP expression, varying from 34.6% to 73.4% relative to the wild type sgRNA (**Figure 5B**). Notably, sgRNAs A4, A19, and A43 significantly reduced sfGFP expression, while A13 and A16 showed minimal inhibitory effects. These findings suggest that the engineered sgRNAs possess varying degrees of binding affinity to dSpRY, enabling differential gene expression repression across a spectrum of efficiencies.

## DISCUSSION

The gene editing technology based on CRISPR/Cas9 has revolutionized existing editing methodologies, propelling the advancement of biological sciences. Multiplex and precise control of gene expression are critical for metabolic regulation in synthetic biology. Co-expressing identical sgRNAs leads to the formation of repetitive sequences and a homogeneity in activity, thereby inducing genetic instability and limiting the breadth of the regulatory scope in the genome editing. To solve the problem of single repetitive sgRNA, sgRNAGen was developed to *de novo* generated diverse sgRNAs based on a decoder Transformer architecture. To assess the effectiveness of sgRNAs generated by sgRNAGen, we synthesized fifty sgRNAs targeting essential genes in *B. subtilis*. The outcomes revealed that a majority of these sgRNAs have activities, demonstrating that sgRNAGen successfully captured the characteristics and learned the distribution of sgRNAs. A series sgRNAs with low similarity and various activities were provided, which could be used in gene editing and metabolic flux redirect. Additionally, to fine-tune metabolic fluxes, we developed a CRISPRi system using our designed sgRNAs. This system features sgRNAs with varying strengths, allowing for regulation of gene expression.

For multiplex gene editing, we successfully knockout two targets with efficiency of 62.5% and 50%, respectively. However, our efforts to concurrently edit multiple genetic sites in *B. subtilis* faced challenges, particularly when attempting to modify more than two sites at once. We hypothesize that this limitation arises due to the increased lethality associated with multiple genomic cuts, coupled with the fact that *B. subtilis* lacks an exogenous recombination system and has limited intrinsic homologous recombination repair capabilities. To overcome these obstacles, two promising strategies can be employed, including enhancing the native homologous recombination repair system and integrating a more efficient homologous recombination system derived from other bacterial species^33^. These approaches could significantly improve repair efficiency during the editing of multiple genomic locations, enabling the simultaneous modification of several sites. Furthermore, the CRISPRi system, which does not require genomic cutting, offers a safer alternative, reducing cell lethality and facilitating the manipulation of multiple targets.

At present, the generated models have a significant challenge lies in assessing the quality of these sequences. In general, two methods were used to evaluate the generated sequences including wet lab verification and computational experiment validation. Here, we mainly used the wet lab verification to access the quality of the generated sgRNAs. While wet lab verification provides precise results, verifying sgRNAs on a large scale is both time-consuming and labor-intensive if it lacks any computational evaluation of the generated sequences. The computational evaluation typically involves using comparative sample distributions and benchmark testing, which depend on assessment datasets. Currently, there is a shortage of high-quality assessment datasets, which is essential for ensuring the accuracy of these evaluations. In addition, the future development of protein-nucleic acid complex structure prediction may assist in the screening and design of RNAs that interact with proteins, potentially reducing the volume of experimental screening required.

Despite the functional application of these sgRNAs in gene editing, their efficiency remains inferior when compared to wild-type sgRNAs. One potential explanation for this discrepancy might be the model focused on the features of sgRNA, neglecting the interaction between Cas9 protein and sgRNA. Furthermore, the performance of the model is significantly influenced by the size and quality of the training dataset. The currently employed crRNA dataset is relatively small, which might lead to incomplete learning. Additionally, the accuracy of sgRNA generation quality assessment based solely on secondary structure may be inadequate. In the future, we are considering the utilization of models that incorporate protein sequences and structures, fully taking into account the impact of Cas9 protein on the entire editing system. Moreover, integrating larger-scale RNA pre-trained models could more effectively capture the characteristics of RNA, thereby enhancing the efficiency and precision of gene editing.

## METHODS

### Details of the sgRNAGen model

We employed one-hot tokenization to train the vocabulary of dataset, an intuitive method for representing textual data. The vocabulary of sgRNAGen encompasses seven tokens: A, T, C, G, and special tokens such as [PAD], [MASK], and [UNK]. By applying one-hot encoding, the input sequences are effectively transformed into numerical vectors, suitable for processing by deep learning models. The hidden size was set as 384. In the sgRNAGen model, positional embedding is incorporated to distinguish the position of each token, similar to the approach used in the GPT2 model. This embedding ensures that the model recognizes the specific placement of tokens within the sequence. These embedding representations are fed into the Transformer layers. We employed a multi-head self-attention mechanism for transformation, with our model comprising 8 layers and 12 heads. To maximize information retention, we introduce residual connections, summing the input to the Transformer layer with the output of the self-attention layer. Following this, the data passes through a new normalization layer and is processed through a two-layer perceptron with a GELU activation function. The final output of the model is the probability of each base at the next position and generates sgRNA sequences by autoregression way from 5’ to 3’, one by one, which the next base has relied on all previously generated bases.

### Training of the sgRNAGen model

For the training of the sgRNAGen model, we utilized 40,000 crRNA sequences sourced from the CRISPRCas-db metagenomics database. The dataset was partitioned into training, testing, and validation subsets in proportions of 0.8, 0.1, and 0.1, respectively. The sgRNAGen model underwent training utilizing the GPT-2 architecture. Before training, the model weights were reinitialized. The batch size was set to 64, allowing for the simultaneous processing of 64 data samples. The model utilizes the Adam optimizer (with β1 = 0.9 and β2 = 0.999) for optimization, with a learning rate of 0.001.

### Strains, Plasmids, and Culture Conditions

All plasmids, primers, and strains utilized in this study are detailed in Tables S1, S2, and S3, respectively. The *E. coli* strain JM109 served as the host for plasmid construction and was cultured in Luria-Bertani (LB) medium at 37 °C. *B. subtilis* was cultured in LB medium at a temperature of 30 °C. To select for target strains, related antibiotics were added to the media: 50 mg/L Kanamycin for *E. coli* and 20 mg/L Kanamycin for *B. subtilis*.

### Plasmid construction

The shuffle plasmid pJOE8999 served as the template for the construction of all editing plasmids. To target the *secY* gene, fifty sgRNAs were introduced into pJOE8999, replacing the wild-type sgRNA, utilizing specifically designed primers. For the development of multiplex gene editing plasmids, we employed overlap extension Polymerase Chain Reaction (PCR) to generate various sgRNA expression cassettes containing distinct sgRNAs. These cassettes were then ligated to the backbone using Gibson Assembly. We utilized the PLCBE plasmid as a template, inactivating its nuclease activity through point mutation techniques and removing the deaminase coding region, thereby obtaining the pXY-dSpRY plasmid. The pXY-dSpRY plasmid serves as a foundational template for CRISPRi related experiments. Different sgRNA sequences are introduced into the system via primers.

### Evaluation of generated sgRNAs activities

Plasmids containing different sgRNAs targeting *SecY* gene were transformed into competent *B. subtilis* 168 cells. After transformation, the cells were spread on LB agar plates containing 20 mg/L kanamycin, with one set supplemented with 0.2% mannose (selective) and the other without mannose (non-selective). The plates were incubated at 37 °C for 8-12 h. The activity of the sgRNAs was assessed by comparing the number of colonies on selective versus non-selective media.

### Gene editing in *B. subtilis*

The editing plasmids were transformed into *B. subtilis*, and the cells were spread on LB agar plates containing 20 mg/L kanamycin and 0.2% mannose, followed by incubation at 30 °C for plate-based editing. For multiplex editing, to enhance the efficiency of editing at multiple loci, single colonies from the plates were picked and inoculated into LB liquid medium supplemented with 20 mg/L kanamycin and 0.2% mannose, followed by incubation for 6-8 h. Subsequently, the cultures were spread onto corresponding LB plates for overnight incubation. The editing efficiency was valided by PCR and Sanger sequencing.

## ASSOCIATED CONTENT

### Supporting information

The Supporting Information is available free of charge at ACS synthetic biology online.

### Author contributions

S.Y.G., Y.-X.H., and Y.X. conceptualized the project. Y.X. designed, built, and implemented deep learning models. Y.X., Z.Y.L., X.W.D., T.D.C., J.L., and L.C.S. carried out the experiments. S.Y.G., Y.-X.H., and Y.X. analyzed the experiments data, wrote the manuscript, and revised the manuscript.

### Notes

The authors declare no competing financial interest.

## Acknowledgements

This work was supported by the National Natural Science Foundation of China (32371489 and 32370095). Part of the experiments was carried out in the Biological & Medical Engineering Core Facilities of the Beijing Institute of Technology.

