## Supplemental information for "Design nonrepetitive and diverse activity single-guide RNA by deep learning"

**Table S1. The plasmids used in this study**

| **Plasmids** | **Description** | **Source** |
| --- | --- | --- |
| pXY-P43-sfGFP | A plasmid expressing sfGFP under the control of the P43 promoter | Laboratory stock |
| p8999-cas9-2N20-HA-21.3 | A genome-scale knockout plasmid harboring two sgRNAs and HA | This study |
| p8999-cas9-2N20-HA-31 | A genome-scale knockout plasmid harboring two sgRNAs and HA | This study |
| p8999-cas9-2N20-HA-27.8 | A genome-scale knockout plasmid harboring two sgRNAs and HA | This study |
| p8999-cas9-2N20-HA-23.3 | A genome-scale knockout plasmid harboring two sgRNAs and HA | This study |
| p8999-cas9-2N20-HA-51.9 | A genome-scale knockout plasmid harboring two sgRNAs and HA | This study |
| p8999-cas9-2N20-HA-169.5 | A genome-scale knockout plasmid harboring two sgRNAs and HA | This study |
| p8999-cas9-bdhA-A3 | A plasmid targeting bdhA by the A3 sgRNA | This study |
| p8999-cas9-bdhA-A4 | A plasmid targeting bdhA by the A4 sgRNA | This study |
| p8999-cas9-bdhA-A7 | A plasmid targeting bdhA by the A7 sgRNA | This study |
| p8999-cas9-bdhA-A15 | A plasmid targeting bdhA by the A15 sgRNA | This study |
| p8999-cas9-bdhA-A37 | A plasmid targeting bdhA by the A37 sgRNA | This study |
| p8999-cas9-bdhA-A38 | A plasmid targeting bdhA by the A38 sgRNA | This study |
| p8999-cas9-bdhA-A44 | A plasmid targeting bdhA by the A44 sgRNA | This study |
| p8999-cas9-bdhA-A49 | A plasmid targeting bdhA by the A49 sgRNA | This study |
| pXY-dSpRY-A4 | Repressing the expression of sfGFP by A4 sgRNA | This study |
| pXY-dSpRY-A5 | Repressing the expression of sfGFP by A5 sgRNA | This study |
| pXY-dSpRY-A7 | Repressing the expression of sfGFP by A7 sgRNA | This study |
| pXY-dSpRY-A19 | Repressing the expression of sfGFP by A19 sgRNA | This study |
| pXY-dSpRY-A21 | Repressing the expression of sfGFP by A21 sgRNA | This study |
| pXY-dSpRY-A25 | Repressing the expression of sfGFP by A25 sgRNA | This study |
| pXY-dSpRY-A27 | Repressing the expression of sfGFP by A27 sgRNA | This study |
| pXY-dSpRY-A29 | Repressing the expression of sfGFP by A29 sgRNA | This study |
| pXY-dSpRY-A33 | Repressing the expression of sfGFP by A33 sgRNA | This study |
| pXY-dSpRY-A38 | Repressing the expression of sfGFP by A38 sgRNA | This study |
| pXY-dSpRY-A43 | Repressing the expression of sfGFP by A43 sgRNA | This study |
| pXY-dSpRY-A44 | Repressing the expression of sfGFP by A44 sgRNA | This study |
| pXY-dSpRY-A46 | Repressing the expression of sfGFP by A46 sgRNA | This study |
| pXY-dSpRY-A49 | Repressing the expression of sfGFP by A49 sgRNA | This study |
| p8999-cas9-nprE-wprA | A plasmid targeting *nprE* and *wprA* | This study |
| p8999-cas9-wprA-epr | A plasmid targeting *wprA* and *epr* | This study |
| p8999-cas9-17.8-19.6 | A plasmid targeting two large fragments | This study |
| p8999-cas9-19.6-23.3 | A plasmid targeting two large fragments | This study |
| p8999-cas9-wprA-epr-nprE | A plasmid targeting *wprA*, *nprE*, and *epr* | This study |
| p8999-cas9-SecY-A1 | Targeting the essential gene *SecY* by A1 sgRNA | This study |
| p8999-cas9-SecY-A2 | Targeting the essential gene *SecY* by A2 sgRNA | This study |
| p8999-cas9-SecY-A3 | Targeting the essential gene *SecY* by A3 sgRNA | This study |
| p8999-cas9-SecY-A4 | Targeting the essential gene *SecY* by A4 sgRNA | This study |
| p8999-cas9-SecY-A5 | Targeting the essential gene *SecY* by A5 sgRNA | This study |
| p8999-cas9-SecY-A6 | Targeting the essential gene *SecY* by A6 sgRNA | This study |
| p8999-cas9-SecY-A7 | Targeting the essential gene *SecY* by A7 sgRNA | This study |
| p8999-cas9-SecY-A8 | Targeting the essential gene *SecY* by A8 sgRNA | This study |
| p8999-cas9-SecY-A9 | Targeting the essential gene *SecY* by A9 sgRNA | This study |
| p8999-cas9-SecY-A10 | Targeting the essential gene *SecY* by A10 sgRNA | This study |
| p8999-cas9-SecY-A11 | Targeting the essential gene *SecY* by A11 sgRNA | This study |
| p8999-cas9-SecY-A12 | Targeting the essential gene *SecY* by A12 sgRNA | This study |
| p8999-cas9-SecY-A13 | Targeting the essential gene *SecY* by A13 sgRNA | This study |
| p8999-cas9-SecY-A14 | Targeting the essential gene *SecY* by A14 sgRNA | This study |
| p8999-cas9-SecY-A15 | Targeting the essential gene *SecY* by A15 sgRNA | This study |
| p8999-cas9-SecY-A16 | Targeting the essential gene *SecY* by A16 sgRNA | This study |
| p8999-cas9-SecY-A17 | Targeting the essential gene *SecY* by A17 sgRNA | This study |
| p8999-cas9-SecY-A18 | Targeting the essential gene *SecY* by A18 sgRNA | This study |
| p8999-cas9-SecY-A19 | Targeting the essential gene *SecY* by A19 sgRNA | This study |
| p8999-cas9-SecY-A20 | Targeting the essential gene *SecY* by A20 sgRNA | This study |
| p8999-cas9-SecY-A21 | Targeting the essential gene *SecY* by A21 sgRNA | This study |
| p8999-cas9-SecY-A22 | Targeting the essential gene *SecY* by A22 sgRNA | This study |
| p8999-cas9-SecY-A23 | Targeting the essential gene *SecY* by A23 sgRNA | This study |
| p8999-cas9-SecY-A24 | Targeting the essential gene *SecY* by A24 sgRNA | This study |
| p8999-cas9-SecY-A25 | Targeting the essential gene *SecY* by A25 sgRNA | This study |
| p8999-cas9-SecY-A26 | Targeting the essential gene *SecY* by A26 sgRNA | This study |
| p8999-cas9-SecY-A27 | Targeting the essential gene *SecY* by A27 sgRNA | This study |
| p8999-cas9-SecY-A28 | Targeting the essential gene *SecY* by A28 sgRNA | This study |
| p8999-cas9-SecY-A29 | Targeting the essential gene *SecY* by A29 sgRNA | This study |
| p8999-cas9-SecY-A30 | Targeting the essential gene *SecY* by A30 sgRNA | This study |
| p8999-cas9-SecY-A31 | Targeting the essential gene *SecY* by A31 sgRNA | This study |
| p8999-cas9-SecY-A32 | Targeting the essential gene *SecY* by A32 sgRNA | This study |
| p8999-cas9-SecY-A33 | Targeting the essential gene *SecY* by A33 sgRNA | This study |
| p8999-cas9-SecY-A34 | Targeting the essential gene *SecY* by A34 sgRNA | This study |
| p8999-cas9-SecY-A35 | Targeting the essential gene *SecY* by A35 sgRNA | This study |
| p8999-cas9-SecY-A36 | Targeting the essential gene *SecY* by A36 sgRNA | This study |
| p8999-cas9-SecY-A37 | Targeting the essential gene *SecY* by A37 sgRNA | This study |
| p8999-cas9-SecY-A38 | Targeting the essential gene *SecY* by A38 sgRNA | This study |
| p8999-cas9-SecY-A39 | Targeting the essential gene *SecY* by A39 sgRNA | This study |
| p8999-cas9-SecY-A40 | Targeting the essential gene *SecY* by A40 sgRNA | This study |
| p8999-cas9-SecY-A41 | Targeting the essential gene *SecY* by A41 sgRNA | This study |
| p8999-cas9-SecY-A42 | Targeting the essential gene *SecY* by A42 sgRNA | This study |
| p8999-cas9-SecY-A43 | Targeting the essential gene *SecY* by A43 sgRNA | This study |
| p8999-cas9-SecY-A44 | Targeting the essential gene *SecY* by A44 sgRNA | This study |
| p8999-cas9-SecY-A45 | Targeting the essential gene *SecY* by A45 sgRNA | This study |
| p8999-cas9-SecY-A46 | Targeting the essential gene *SecY* by A46 sgRNA | This study |
| p8999-cas9-SecY-A47 | Targeting the essential gene *SecY* by A47 sgRNA | This study |
| p8999-cas9-SecY-A48 | Targeting the essential gene *SecY* by A48 sgRNA | This study |
| p8999-cas9-SecY-A49 | Targeting the essential gene *SecY* by A49 sgRNA | This study |
| p8999-cas9-SecY-A50 | Targeting the essential gene *SecY* by A50 sgRNA | This study |

**Table S2. The Primers used in this study.**

| **Primers** | **Sequences (5’-3’)** |
| --- | --- |
| A1-F | AAGGCAAAATAATGCTAGTCCGTTCCCAACTTGAAAAAGTGACTCCATCTGGATTTGTTCAGAAC |
| A1-R | GGACTAGCATTATTTTGCCTTGCTATTTCTAGCTGCAAAACCTCGGTAAACTTCGGTACCAC |
| A2-F | AAGTTAAAACACGGCACCGCCACTTTTAACCTAACAAGGTGACTCCATCTGGATTTGTTCAGAAC |
| A2-R | GGCGGTGCCGTGTTTTAACTTGCACTATTGAACCCTAAAACCTCGGTAAACTTCGGTACCAC |
| A3-F | ACGTTAAGACAAGGCATAGCCGCAACAGACTGGACAGGGTGACTCCATCTGGATTTGTTCAGAAC |
| A3-R | GGCTATGCCTTGTCTTAACGTACTATAATTAGTCCTAAAACCTCGGTAAACTTCGGTACCAC |
| A4-F | ACGTTAAGACAAGGCGTAGCCACGACAGACTGGACACGGTGACTCCATCTGGATTTGTTCAGAAC |
| A4-R | GGCTACGCCTTGTCTTAACGTACTATTAGTAGTGCTAAAACCTCGGTAAACTTCGGTACCAC |
| A5-F | TGGTTGAGATACGTTATGATATCTATCTACTTGAGAAAGTCACTCCATCTGGATTTGTTCAGAAC |
| A5-R | TATCATAACGTATCTCAACCAGCCATACATAACTCTGAAACCTCGGTAAACTTCGGTACCAC |
| A6-F | AAGTTAAAATAAGGCTAGACCGTTATCAACTAGAAATAGTGACTCCATCTGGATTTGTTCAGAAC |
| A6-R | GGTCTAGCCTTATTTTAACTTGTGATTTCTCACTCTAAAACCTCGGTAAACTTCGGTACCAC |
| A7-F | AAGTTAAGATATGATTGCTCCCTAGAAGACTTGATAAGGTGACTCCATCTGGATTTGTTCAGAAC |
| A7-R | GGAGCAATCATATCTTAACTTGCTTAGCAAAGCGCTAAAATCTCGGTAAACTTCGGTACCAC |
| A8-F | AAGTAGATATAAGGCCAGTCCGTTAGCAACTTGAAAAAGTCACTCCATCTGGATTTGTTCAGAAC |
| A8-R | GGACTGGCCTTATATCTACTTACTCTGTCGAGTTCAGATACCTCGGTAAACTTCGGTACCAC |
| A9-F | AAGTTAAGATATGATTGCTCCCTAGAAGACTTGATAAGGTGACTCCATCTGGATTTGTTCAGAAC |
| A9-R | GGAGCAATCATATCTTAACTTGCTTAGCAAAGCGCTAAAATCTCGGTAAACTTCGGTACCAC |
| A10-F | GAGTTAAAATAAGGCAAGTCCATTAACCACTTTAAAAAGTAACTCCATCTGGATTTGTTCAGAAC |
| A10-R | GGACTTGCCTTATTTTAACTCGCCATTTCTAGCGCTAAAACCTCGGTAAACTTCGGTACCAC |
| A11-F | ACGTTAAAATAAGGCTAGGCCGTGATATACTTAAAAAAGTAACTCCATCTGGATTTGTTCAGAAC |
| A11-R | GGCCTAGCCTTATTTTAACGTGCTATATATAGCTCTAAAACCTCGGTAAACTTCGGTACCAC |
| A12-F | ACGTTGAAATAAGACTAGGCCGAGACATACTTCAAAAAGTAACTCCATCTGGATTTGTTCAGAAC |
| A12-R | GGCCTAGTCTTATTTCAACGTGCTATATTTAGCTCTAAAACCTCGGTAAACTTCGGTACCAC |
| A13-F | AAGTTAAGATAAGCCTACTCTATTATTTACTTGAAGAAGTGACTCCATCTGGATTTGTTCAGAAC |
| A13-R | AGAGTAGGCTTATCTTAACTTGCTACTTACAGCTCTAAAACCTCGGTAAACTTCGGTACCAC |
| A14-F | TAGTTAAGATAAGCCTAGTCGATTAATTGCTTGAAGAAGTGACTCCATCTGGATTTGTTCAGAAC |
| A14-R | CGACTAGGCTTATCTTAACTAGCTACTTCCAGCTCTAAAACCTCGGTAAACTTCGGTACCAC |
| A15-F | CCGTTAAGATAAGCCCGGTCGATTTATTGCTTGAAGAAGTGACTCCATCTGGATTTGTTCAGAAC |
| A15-R | CGACCGGGCTTATCTTAACGGGCTACGTCCAGCTCTAAAACCTCGGTAAACTTCGGTACCAC |
| A16-F | AGGTTAGAATAAGGTTAGTAATTTATCTACTTCAAGGAGTAACTCCATCTGGATTTGTTCAGAAC |
| A16-R | TTACTAACCTTATTCTAACCTGCTATATTTAGCTCTAAAACCTCGGTAAACTTCGGTACCAC |
| A17-F | ACGTTAAAATAAGCCTAATCTTTGATATACTTTAAAAAGTAACTCCATCTGGATTTGTTCAGAAC |
| A17-R | AGATTAGGCTTATTTTAACGTGCTATAGGTAGCTCTAAAACCTCGGTAAACTTCGGTACCAC |
| A18-F | GAGTTAAAGCAAGACTACTCCGATATCTACTTGCAAGAGTGACTCCATCTGGATTTGTTCAGAAC |
| A18-R | GGAGTAGTCTTGCTTTAACTCACTAATTTTAGCACTAAAACCTCGGTAAACTTCGGTACCAC |
| A19-F | AAGTTAAAATAAGACTAACTCGGAATCGACTGTAAAAAGTGACTCCATCTGGATTTGTTCAGAAC |
| A19-R | GAGTTAGTCTTATTTTAACTTGCTATGGTTAGCTCTAAAACCTCGGTAAACTTCGGTACCAC |
| A20-F | AAGTTAAAATAAGAGTAACTTAGAATCGACTGTAAAAAGTGACTCCATCTGGATTTGTTCAGAAC |
| A20-R | AAGTTACTCTTATTTTAACTTGCTATAGTTAGCTCTAAAACCTCGGTAAACTTCGGTACCAC |
| A21-F | AAGTTGAAGTACGGGTACGCCGCATACTACTTGAAAAAGGGACTCCATCTGGATTTGTTCAGAAC |
| A21-R | GGCGTACCCGTACTTCAACTTCCTATGCATAGATCTGAAATCTCGGTAAACTTCGGTACCAC |
| A22-F | TAGTTAAGACAAGGGTAGACCGTTACGCACTGGAATAAGTGACTCCATCTGGATTTGTTCAGAAC |
| A22-R | GGTCTACCCTTGTCTTAACTAGCTATAATCAGCTCTAAAACCTCGGTAAACTTCGGTACCAC |
| A23-F | ATGTTAAAATAAGATTGACCGGTAATATACTTAACGAAGTGACTCCATCTGGATTTGTTCAGAAC |
| A23-R | CGGTCAATCTTATTTTAACATGCTACATCTAGCTCTAAAACCTCGGTAAACTTCGGTACCAC |
| A24-F | ATGTTAGAATAAGATTTACCGATAATATACTTAACGAAGTGACTCCATCTGGATTTGTTCAGAAC |
| A24-R | CGGTAAATCTTATTCTAACATGCTACCTCTAGCTCTAAAACCTCGGTAAACTTCGGTACCAC |
| A25-F | TACGTTAGAACATGGGCGCCCCTTTATTCACTGTAAAAAGTGACTCCATCTGGATTTGTTCAGAAC |
| A25-R | GGGCGCCCATGTTCTAACGTACTATTGCTAGCTCTAAAACCTCGGTAAACTTCGGTACCAC |
| A26-F | TAAGTTAAAACATGGGCGATCCTTAATTCACTGAAGAAAGTGACTCCATCTGGATTTGTTCAGAAC |
| A26-R | GATCGCCCATGTTTTAACTTACTATTGCTAGCACTAAAACCTCGGTAAACTTCGGTACCAC |
| A27-F | TAGGTTGAAACATGGTCGATCCTTAATTAACTGAAGAAAGTGACTCCATCTGGATTTGTTCAGAAC |
| A27-R | GATCGACCATGTTTCAACCTACTATCGCTAGCACTAAAACCTCGGTAAACTTCGGTACCAC |
| A28-F | CCAGTTGAAGCACGACTCTTCCCTTATATACTTGAGAAAGTGACTCCATCTGGATTTGTTCAGAAC |
| A28-R | GAAGAGTCGTGCTTCAACTGGCTAATAGTAACTCTAAAGCCTCGGTAAACTTCGGTACCAC |
| A29-F | CTTGTTGAAATACGGCGTCTCCGATTTAGACTTGAGGAAGTGACTCCATCTGGATTTGTTCAGAAC |
| A29-R | GAGACGCCGTATTTCAACAAGCTGTGGTTAGCACTAAAACCTCGGTAAACTTCGGTACCAC |
| A30-F | AAAGTTAAAATAAGGATATGCCACATTCAACTGTCAAGAGTAACTCCATCTGGATTTGTTCAGAAC |
| A30-R | GCATATCCTTATTTTAACTTTCTATGTGTAGTGCTAAAACCTCGGTAAACTTCGGTACCAC |
| A31-F | CAAGTTGAAATATGCCCAAACCGTTCCACGCTTCTAAGAGTTACTCCATCTGGATTTGTTCAGAAC |
| A31-R | GTTTGGGCATATTTCAACTTGCTATCTCTAGCTCAGAAACCTCGGTAAACTTCGGTACCAC |
| A32-F | AAGGTTGAAATACGTTTGATCCATTATATACTTGAAGAAGTCACTCCATCTGGATTTGTTCAGAAC |
| A32-R | GATCAAACGTATTTCAACCTTCCAAATGTGGATCTAAAACCTCGGTAAACTTCGGTACCAC |
| A33-F | CATGTTAAAATACGGGCTTGCCGTCTACGACCTAAAGAGGTAACTCCATCTGGATTTGTTCAGAAC |
| A33-R | GCAAGCCCGTATTTTAACATGCTAATTCAAACTCTAAAATCTCGGTAAACTTCGGTACCAC |
| A34-F | CAGGTTAGGACAAGATTCGCAAGTTCATCACTTAAAGAAGTGACTCCATCTGGATTTGTTCAGAAC |
| A34-R | TGCGAATCTTGTCCTAACCTGCTAAAGGTAGCTCTAAAACCTCGGTAAACTTCGGTACCAC |
| A35-F | TATGTTGAGACAAGAATCCTCTGATAAGTACTTGATAAGGTAACTCCATCTGGATTTGTTCAGAAC |
| A35-R | GAGGATTCTTGTCTCAACATACCAATACTAGCCCTAAAACCTCGGTAAACTTCGGTACCAC |
| A36-F | TCCGTTGAGACAAGAATTCTCTGATAAGTATCTGATAAGGTTACTCCATCTGGATTTGTTCAGAAC |
| A36-R | GAGAATTCTTGTCTCAACGGACCAAGACTAGCCCTAAAACCTCGGTAAACTTCGGTACCAC |
| A37-F | CATGTTAATACAAGCCCACACATTGATCTACTTGAAAAAGTTACTCCATCTGGATTTGTTCAGAAC |
| A37-R | GTGTGGGCTTGTATTAACATGCTATTTGAAGCTCTAATACCTCGGTAAACTTCGGTACCAC |
| A38-F | CAAGTCAAAACAAGGCTAGACCAATATAAACTTTTTAAAGTTACTCCATCTGGATTTGTTCAGAAC |
| A38-R | GTCTAGCCTTGTTTTGACTTGCTACATGTAGCTCTAAAACCTCGGTAAACTTCGGTACCAC |
| A39-F | CCAGTTAGGATAAGACTATTTCGTTATACACTTCACAAAGTAACTCCATCTGGATTTGTTCAGAAC |
| A39-R | AAATAGTCTTATCCTAACTGGCTATATTTAGCTCTAGAACCTCGGTAAACTTCGGTACCAC |
| A40-F | CCGGTTAGGATACGAATATTGCGTTACACACTTCCAAAAGTAACTCCATCTGGATTTGTTCAGAAC |
| A40-R | CAATATTCGTATCCTAACCGGCTATTTTTAGCTCTAGAACCTCGGTAAACTTCGGTACCAC |
| A41-F | CCGGTTAGAATACGTAAATTGCGGTACACACTTCCAAAAGTAACTCCATCTGGATTTGTTCAGAAC |
| A41-R | CAATTTACGTATTCTAACCGGCTAGCATTAGCACTAGAACCTCGGTAAACTTCGGTACCAC |
| A42-F | CAAGTTAAAATAAGGTTAGCACATATTCCACTTAACGAAGTGACTCCATCTGGATTTGTTCAGAAC |
| A42-R | TGCTAACCTTATTTTAACTTGCTACAAGTAGCTCTAAAACCTCGGTAAACTTCGGTACCAC |
| A43-F | CGGGTTAAAATAAGGCTTTGCCTAATTCTACTTTTGAAAGTGACTCCATCTGGATTTGTTCAGAAC |
| A43-R | GCAAAGCCTTATTTTAACCCGCTATTCTTAGCTCTAAAACCTCGGTAAACTTCGGTACCAC |
| A44-F | CACGTTGAAATAAGGCATGTTCATTCTCCACTTTTTTAAGTGACTCCATCTGGATTTGTTCAGAAC |
| A44-R | AACATGCCTTATTTCAACGTGCTATAACTAGCTCTAAAACCTCGGTAAACTTCGGTACCAC |
| A45-F | CAAGTTAAAATATGGCAACTTTTATATCTACTTGGCAGAGTGACTCCATCTGGATTTGTTCAGAAC |
| A45-R | AAGTTGCCATATTTTAACTTGCTGTTAGCAGCTCTAAAACCTCGGTAAACTTCGGTACCAC |
| A46-F | CAAGTTAAAATAGGGCAACTTTTATATCCACTTGACAGAGTGACTCCATCTGGATTTGTTCAGAAC |
| A46-R | AAGTTGCCCTATTTTAACTTGCTGTTCGCAGCTCTAAAACCTCGGTAAACTTCGGTACCAC |
| A47-F | CAAGTTAAAATAGGCCAACTTTCGTATCCACTTGACAGAGTGACTCCATCTGGATTTGTTCAGAAC |
| A47-R | AAGTTGGCCTATTTTAACTTGCTGTTGGCAGCTCTAAAATCTCGGTAAACTTCGGTACCAC |
| A48-F | CAAGTTAAAATAAGCCAACTATCGTATCCACTTGTCAGAGTGACTCCATCTGGATTTGTTCAGAAC |
| A48-R | TAGTTGGCTTATTTTAACTTGCTGTGTTCAGCTCTAAAATCTCGGTAAACTTCGGTACCAC |
| A49-F | CAAGTTAAGATAAGGCTGGTACGAATTACACTTTCTAAAGTGACTCCATCTGGATTTGTTCAGAAC |
| A49-R | TACCAGCCTTATCTTAACTTGCCATAACTAGCTCTAAAACCTCGGTAAACTTCGGTACCAC |
| A50-F | CAAGTTAGAATAAGCATTGCATATCACCAACTTGAAAAAGTGACTCCATCTGGATTTGTTCAGAAC |
| A50-R | TGCAATGCTTATTCTAACTTGCCATTTTTAGCGCTAAAACCTCGGTAAACTTCGGTACCAC |
| B3-F | GTACGTTAAGACAAGGCATAGCCGCAACAGACTGGACAGGGTGACTCCATCTGGATTTGTTCAGAAC |
| B3-R | CTATGCCTTGTCTTAACGTACTATAATTAGTCCTAAAACCCTGTACGTTCTGCAATCTCC |
| B4-F | TACGTTAAGACAAGGCGTAGCCACGACAGACTGGACACGGTGACTCCATCTGGATTTGTTCAGAAC |
| B4-R | CTACGCCTTGTCTTAACGTACTATTAGTAGTGCTAAAACCCTGTACGTTCTGCAATCTCC |
| B7-F | GCAAGTTAAGATATGATTGCTCCCTAGAAGACTTGATAAGGTGACTCCATCTGGATTTGTTCAGAAC |
| B7-R | GCAATCATATCTTAACTTGCTTAGCAAAGCGCTAAAATCCTGTACGTTCTGCAATCTCC |
| B15-F | CGACCGGGCTTATCTTAACGGGCTACGTCCAGCTCTAAAACCCTGTACGTTCTGCAATCTCC |
| B15-R | CGTTAAGATAAGCCCGGTCGATTTATTGCTTGAAGAAGTGACTCCATCTGGATTTGTTCAGAAC |
| B37-F | TGTTAATACAAGCCCACACATTGATCTACTTGAAAAAGTTACTCCATCTGGATTTGTTCAGAAC |
| B37-R | AATGTGTGGGCTTGTATTAACATGCTATTTGAAGCTCTAATACCCTGTACGTTCTGCAATCTC |
| B38-F | AGTCAAAACAAGGCTAGACCAATATAAACTTTTTAAAGTTACTCCATCTGGATTTGTTCAGAAC |
| B38-R | GGTCTAGCCTTGTTTTGACTTGCTACATGTAGCTCTAAAACCCTGTACGTTCTGCAATCTC |
| B44-F | CGTTGAAATAAGGCATGTTCATTCTCCACTTTTTTAAGTGACTCCATCTGGATTTGTTCAGAAC |
| B44-R | GAACATGCCTTATTTCAACGTGCTATAACTAGCTCTAAAACCCTGTACGTTCTGCAATCTC |
| B49-F | GCAAGTTAAGATAAGGCTGGTACGAATTACACTTTCTAAAGTGACTCCATCTGGATTTGTTCAGAAC |
| B49-R | CCAGCCTTATCTTAACTTGCCATAACTAGCTCTAAAACCCTGTACGTTCTGCAATCTC |
| Δ21.3-N20-1 | gacaaattggtatctgacatgttttagagctagaaatagcaagttaaaataagg |
| Δ21.3-N20-2 | atgtcagataccaatttgtccgtaggtacattttactcaattctctaatc |
| Δ21.3-N20-3 | CACAAGTTAAAATAAGGCTAGACCGTTATCAACTAGAAATAGTGTTTTTACTCCATCTGGATTTGTTCAGAACGCTCGG |
| Δ21.3-N20-4 | CGGTCTAGCCTTATTTTAACTTGTGATTTCTCACTCTAAAACTTCCTGGGTAGAGGTGTATTCGTAGGTACAT |
| Δ21.3-HA-1 | cttctcccccattacatcacttgtacatgtcgatacctctatgctg |
| Δ21.3-HA-2 | gtcttctatgtcgtaccttagccaaatcgctattca |
| Δ21.3-HA-3 | ggctaaggtacgacatagaagacatgtcaaagc |
| Δ21.3-HA-4 | ggaggtgactgaagtataccgaatacggagaagaccatggtctaca |
| Δ21.3-Test-1 | ttgtaccgatgttgacgttgat |
| Δ21.3-Test-2 | acacgtaaatgaattcattaccgat |
| Δ21.3-Test-3 | ttacttggtataatggagtcggga |
| Δ21.3-Test-4 | ttcctaagcggaaatagatggt |
| Δ31-N20-1 | ccgatccagtcatacagatcgttttagagctagaaatagcaagttaaaataagg |
| Δ31-N20-2 | gatctgtatgactggatcggcgtaggtacattttactcaattctctaatc |
| Δ31-N20-3 | CACAAGTTAAAATAAGGCTAGACCGTTATCAACTAGAAATAGTGTTTTTACTCCATCTGGATTTGTTCAGAACGCTCGG |
| Δ31-N20-4 | CGGTCTAGCCTTATTTTAACTTGTGATTTCTCACTCTAAAACCTCTGTAGTTGTTCAGTTGACGTAGG |
| Δ31-HA-1 | cttctcccccattacatcacttctgtttgtcgggtgaaacg |
| Δ31-HA-2 | actggatatgattcgtcaagcggataaattcttgc |
| Δ31-HA-3 | gcttgacgaatcatatccagtttcatccactcgg |
| Δ31-HA-4 | gaggtgactgaagtataccgaaacatacctgatatctgcatgaatgc |
| Δ31-Test-1 | tcaatcgaaaatgctgtcaccat |
| Δ31-Test-2 | tcttgatgatagccaagctgc |
| Δ31-Test-3 | aactgatgatcgtaattggcca |
| Δ31-Test-4 | acatgcataggaaagatttccg |
| Δ27.8-N20-1 | gggtcttcctgaaatcctgagttttagagctagaaatagcaagttaaaataaggc |
| Δ27.8-N20-2 | tcaggatttcaggaagaccccgtaggtacattttactcaattctctaatca |
| Δ27.8-N20-3 | CACAAGTTAAAATAAGGCTAGACCGTTATCAACTAGAAATAGTGTTTTTACTCCATCTGGATTTGTTCAGAACGCTCGGTTGCCG |
| Δ27.8-N20-4 | CGGTCTAGCCTTATTTTAACTTGTGATTTCTCACTCTAAAACTCCCTGTTCTGCTTGTCTATCGTAGGTACAT |
| Δ27.8-HA-1 | cttctcccccattacatcactaattggccttttcctgtacc |
| Δ27.8-HA-2 | tttgtctcagctgaagcagg |
| Δ27.8-HA-3 | cctgcttcagctgagacaaagtcaattttcgcggactctt |
| Δ27.8-HA-4 | aggaggtgactgaagtataccgaatgacctctgaacagatcggc |
| Δ27.8-Test-1 | tgtccgcatctatattcagaagc |
| Δ27.8-Test-2 | aagctattatctgtgccgtgc |
| Δ27.8-Test-3 | tctgtcagctcacggtattct |
| Δ27.8-Test-4 | atgcaggctgatgaactagatat |
| Δ23.3-N20-1 | gaatttagctccgccgataagttttagagctagaaatagcaagttaaaataagg |
| Δ23.3-N20-2 | ttatcggcggagctaaattccgtaggtacattttactcaattctctaatc |
| Δ23.3-N20-3 | CACAAGTTAAAATAAGGCTAGACCGTTATCAACTAGAAATAGTGTTTTTACTCCATCTGGATTTGTTCAGAACGCTCGGTTGCCG |
| Δ23.3-N20-4 | CGGTCTAGCCTTATTTTAACTTGTGATTTCTCACTCTAAAACGCAAACAACTGCCGATCAAGCGTAGGTAC |
| Δ23.3-HA-1 | cttctcccccattacatcactagaacttcagcgattctcgaa |
| Δ23.3-HA-2 | catgagcgagttatgtacagcctgtattcattgct |
| Δ23.3-HA-3 | gctgtacataactcgctcatgttgatgtct |
| Δ23.3-HA-4 | gaggtgactgaagtataccgaattcttgacaccagcttatgcaa |
| Δ23.3-Test-1 | tggcaccatttatatttggtctct |
| Δ23.3-Test-2 | gtgcagacaagaacagtagaact |
| Δ23.3-Test-3 | tatatggaagaacgatcacagcg |
| Δ23.3-Test-4 | tatctataatggcgctcagcac |
| Δ51.9-N20-1 | attgtcattccgatcatggcgttttagagctagaaatagcaagttaaaataagg |
| Δ51.9-N20-2 | gccatgatcggaatgacaatcgtaggtacattttactcaattctctaatc |
| Δ51.9-N20-3 | GTGAGAAATCACAAGTTAAAATAAGGCTAGACCGTTATCAACTAGAAATAGTGTTTTTACTCCATCTGGATTTGTTCAGAACGCTCGG |
| Δ51.9-N20-4 | GCCTTATTTTAACTTGTGATTTCTCACTCTAAAACTCAGCTGTGTCAATGATGTGCGTAGGTAC |
| Δ51.9-HA-1 | ttctcccccattacatcactacgccaataatgcttcaatcg |
| Δ51.9-HA-2 | ttcggctccatttatattcatggctttcgcaaacc |
| Δ51.9-HA-3 | tgaatataaatggagccgaaaatcaccatag |
| Δ51.9-HA-4 | gaggtgactgaagtataccgaatgtagacaaggttgaagaggacg |
| Δ51.9-Test-1 | ataaagaaagctccagtggcaa |
| Δ51.9-Test-2 | tactacatgtcgatggcggc |
| Δ51.9-Test-3 | ttcttggaatacgatgatgaccg |
| Δ51.9-Test-4 | atgccttgctcaggagtaaac |
| C1 | TTAGAAATGGGCGTGAAAAAAAGCGCGCGATTATGTAAAATATAGAGTGATAGCGGTACCATTATAGGTAAGAGAGGAATG |
| C2 | ACTTACTCTGTCGAGTTCAGATACCTGCCTTAACGTCTTTTACCGTGTACATTCCTCTCTTACCTATAATGG |
| C3 | GTATCTGAACTCGACAGAGTAAGTAGATATAAGGCCAGTCCGTTAGCAACTTGAAAAAGTCTGATTCAAGC |
| C4 | AACTGCGTTTGACCAAAACAAACAAAAAACCCAGCTCATTGAGCTGGGTTTAAGCTTGCTTGAATCAGACTTTTTCAAGTT |
| C5 | TGTTTTGGTCAAACGCAGTTAGCTTGGCCGAATTCGTCGACTCTAGACTGCAGATCGTCGAACGGCAGATCAGAATTT |
| C6 | TTACCATGAAAAAGAGCCCGCAGTGTAATGAGCAGGCTCTTTTTTTATTACAAAATTCTGATCTGCCGTTCGA |
| C7 | GCGGGCTCTTTTTCATGGTAAAAAACGGCCTCTCGAAATAGAGGGTTGACACTCTTTTGAGAA |
| C8 | TTTATCACCTCCTTTCTGATAATATAACATATTCTCAAAAGAGTGTCAACCCT |
| C9 | TGTTATATTATCAGAAAGGAGGTGATAAAACGGAAAAAGCAGTACAGAAAGATGTAGATGTAGAAATACAAGGTTACAT |
| C10 | CGGCCGTTCCACTTCTTCAAGTTGATTACGGACGGGCCTTAATGTAACCTTGTATTTCTACATCTAC |
| C11 | TGAAGAAGTGGAACGGCCGGATGTCTTAAAAAAGACGTCCGGCTTTTCTTTTTGTATCCATGATCGTAGTGCAAGTACA |
| C12 | ATCAACGTTAATAAGACGTTGTCTTTGCAATAAAAAAACCGTCCGGCGTACGCCGGGCGGTTTGTGTACTTGCACTACGATCAT |
| C13 | AGACAACGTCTTATTAACGTTGATATAATTTAAATTTTATTTGACAAAAATGGGCTCGTGTTGTACAATAAATGT |
| C14 | ATTTTAACTTGCTATTTCTAGCTCTAAAACTGCAACGCTCGGATCTTTTTACATTTATTGTACAACACGAGCC |
| C15 | GCTAGAAATAGCAAGTTAAAATAAGGCTAGTCCGTTATCAACTTGAAAAAGTGGGCAACAGAATTTGCCTG |
| C16 | TCACTTCTGAGTTCGGCATGGGGTCAGGTGGGACCACCGCGCTACTGCCGCCAGGCAAATTCTGTTGCCCA |
| C17 | CATGCCGAACTCAGAAGTGAAACGCCGTAGCGCCGATGGTAGTGTGGGATCTCCCCATGCGAGAGTAGGGAA |
| C18 | GTCTTTCGACTGAGCCTTTCGTTTTATTTGATGCCTGGCAGTTCCCTACTCTCGCATGGG |
| C1-F | TAACTTGCTTAGCAAAGCGCTAAAATCTTTTGAGAATATGTTATATACATTTATTGTACAACAC |
| C1-R | GCGCTTTGCTAAGCAAGTTAAGATATGATTGCTCCCTAGAAGACTTGATAAGGTGGGCAACAGAATTTGCCTGG |
| C2-F | CGGAGTAGTCTTGCTTTAACTCACTAATTTTAGCACTAAAACCTTTTGAGAATATGTTATATACATTTATTGTACAACAC |
| C2-R | GTTTTAGTGCTAAAATTAGTGAGTTAAAGCAAGACTACTCCGATATCTACTTGCAAGAGTG |
| C3-F | CCGGTCAATCTTATTTTAACATGCTACATCTAGCTCTAAAACCTTTTGAGAATATGTTATATACATTTATTGTACAACAC |
| C3-R | GCATGTTAAAATAAGATTGACCGGTAATATACTTAACGAAGTGGGCAACAGAATTTGCCTGG |
| C4-F | CGCCCATGTTCTAACGTACTATTGCTAGCTCTAAAACCTTTTGAGAATATGTTATATACATTTATTGTACAACAC |
| C4-R | AGTACGTTAGAACATGGGCGCCCCTTTATTCACTGTAAAAAGTGGGCAACAGAATTTGCCTGG |
| C5-F | TTCAACCTACTATCGCTAGCACTAAAACCTTTTGAGAATATGTTATATACATTTATTGTACAACAC |
| C5-R | GTGCTAGCGATAGTAGGTTGAAACATGGTCGATCCTTAATTAACTGAAGAAAGTGGGCAACAGAATTTGCCTGG |
| C6-F | AATCGGAGACGCCGTATTTCAACAAGCTGTGGTTAGCACTAAAACCTTTTGAGAATATGTTATATACATTTATTGTACAACAC |
| C6-R | AATACGGCGTCTCCGATTTAGACTTGAGGAAGTGGGCAACAGAATTTGCCTGG |
| C7-F | CGGCAAGCCCGTATTTTAACATGCTAATTCAAACTCTAAAATCTTTTGAGAATATGTTATATACATTTATTGTACAACAC |
| C7-R | AAAATACGGGCTTGCCGTCTACGACCTAAAGAGGTAGGCAACAGAATTTGCCTGG |
| C8-F | GGTCTAGCCTTGTTTTGACTTGCTACATGTAGCTCTAAAACCTTTTGAGAATATGTTATATACATTTATTGTACAACAC |
| C8-R | AAGTCAAAACAAGGCTAGACCAATATAAACTTTTTAAAGTTGGCAACAGAATTTGCCTGG |
| C9-F | TTCAACGTGCTATAACTAGCTCTAAAACCTTTTGAGAATATGTTATATACATTTATTGTACAACAC |
| C9-R | AGAGCTAGTTATAGCACGTTGAAATAAGGCATGTTCATTCTCCACTTTTTTAAGTGGGCAACAGAATTTGCCTGG |
| C10-F | CGTACCAGCCTTATCTTAACTTGCCATAACTAGCTCTAAAACCTTTTGAGAATATGTTATATACATTTATTGTACAACAC |
| C10-R | CAAGTTAAGATAAGGCTGGTACGAATTACACTTTCTAAAGTGGGCAACAGAATTTGCCTGG |
| C11-F | GGCTACGCCTTGTCTTAACGTACTATTAGTAGTGCTAAAACCTTTTGAGAATATGTTATATACATTTATTGTACAACAC |
| C11-R | GTTAAGACAAGGCGTAGCCACGACAGACTGGACACGGTGGGCAACAGAATTTGCCTGG |
| C12-F | AGAGTAGGCTTATCTTAACTTGCTACTTACAGCTCTAAAACCTTTTGAGAATATGTTATATACATTTATTGTACAACAC |
| C12-R | GTAGCAAGTTAAGATAAGCCTACTCTATTATTTACTTGAAGAAGTGGGCAACAGAATTTGCCTGG |
| C13-F | CCGAGTTAGTCTTATTTTAACTTGCTATGGTTAGCTCTAAAACCTTTTGAGAATATGTTATATACATTTATTGTACAACAC |
| C13-R | AGCAAGTTAAAATAAGACTAACTCGGAATCGACTGTAAAAAGTGGGCAACAGAATTTGCCTGG |
| C14-F | CGGTAAATCTTATTCTAACATGCTACCTCTAGCTCTAAAACCTTTTGAGAATATGTTATATACATTTATTGTACAACAC |
| C14-R | GGTAGCATGTTAGAATAAGATTTACCGATAATATACTTAACGAAGTGGGCAACAGAATTTGCCTGG |
| C15-F | GGATCAAACGTATTTCAACCTTCCAAATGTGGATCTAAAACCTTTTGAGAATATGTTATATACATTTATTGTACAACAC |
| C15-R | GGAAGGTTGAAATACGTTTGATCCATTATATACTTGAAGAAGTCGGCAACAGAATTTGCCTGG |
| C16-F | GCAATGCTTATTCTAACTTGCCATTTTTAGCGCTAAAACTTATCAGAAAGGAGGTGATAACATTTATTGTA |
| C16-R | TGGCAAGTTAGAATAAGCATTGCATATCACCAACTTGAAAAAGTGGGCAACAGAATTTGCCTGG |
| C17-F | CCTTATTCTAACCTGCTATATTTAGCTCTAAAACCTTTTGAGAATATGTTATATACATTTATTGTACAACAC |
| C17-R | GAGCTAAATATAGCAGGTTAGAATAAGGTTAGTAATTTATCTACTTCAAGGAGTAGGCAACAGAATTTGCCTGG |
| C18-F | TGTGGGCTTGTATTAACATGCTATTTGAAGCTCTAATACTTATCAGAAAGGAGGTGATAACATTTATTGTA |
| C18-R | GCATGTTAATACAAGCCCACACATTGATCTACTTGAAAAAGTTGGCAACAGAATTTGCCTGG |
| C19-F | AACCCGCTATTCTTAGCTCTAAAACTTATCAGAAAGGAGGTGATAACATTTATTGTA |
| C19-R | AGAGCTAAGAATAGCGGGTTAAAATAAGGCTTTGCCTAATTCTACTTTTGAAAGTGGGCAACAGAATTTGCCTGG |
| C20-F | TTAACTTGCTGTTCGCAGCTCTAAAACTTATCAGAAAGGAGGTGATAACATTTATTGTA |
| C20-R | GCTGCGAACAGCAAGTTAAAATAGGGCAACTTTTATATCCACTTGACAGAGTGGGCAACAGAATTTGCCTGG |
| C31-F | GTGCGGAATTACTCACATCAACATTTATTGTACAACACGAGCC |
| C31-R | TGATGTGAGTAATTCCGCACGTTTTAGAGCTAGAAATAGCAAGTTAAAAT |
| C32-F | AAAGTTATAGGAGTGGTAATACATTTATTGTACAACACGAGCC |
| C32-R | ATTACCACTCCTATAACTTTGTTTTAGAGCTAGAAATAGCAAGTTAAAAT |
| C33-F | TGAAGGCAGCAAGATGGCATACATTTATTGTACAACACGAGCC |
| C33-R | ATGCCATCTTGCTGCCTTCAGTTTTAGAGCTAGAAATAGCAAGTTAAAAT |

**Table S3. The strains used in this study.**

| **Strains** | **Description** | **Source** |
| --- | --- | --- |
| *E. coli* JM109 | *recA1*, *endA1*, *thi*, *gyrA96*, *supE44*, *hsdR17*∆ (*lac-proAB*) /F’[*traD36*,*proAB*^+^, *lacІ^q^*, *lacZ∆ M15*] | Laboratory stock |
| *B. subtilis* | Wild-type *Bacillus subtilis* 168 | Laboratory stock |
| BS-P43-sfGFP | Wild-type *B. subtilis* carrying the plasmid PXY-P43-sfGFP | Laboratory stock |
| p8999-cas9-2N20-HA-21.3 | Wild-type *B. subtilis* carrying the plasmid p8999-cas9-2N20-HA-21.3 | This study |
| p8999-cas9-2N20-HA-31 | Wild-type *B. subtilis* carrying the plasmid p8999-cas9-2N20-HA-31 | This study |
| p8999-cas9-2N20-HA-27.8 | Wild-type *B. subtilis* carrying the plasmid p8999-cas9-2N20-HA-27.8 | This study |
| p8999-cas9-2N20-HA-23.3 | Wild-type *B. subtilis* carrying the plasmid p8999-cas9-2N20-HA-23.3 | This study |
| p8999-cas9-2N20-HA-51.9 | Wild-type *B. subtilis* carrying the plasmid p8999-cas9-2N20-HA-51.9 | This study |
| p8999-cas9-2N20-HA-169.5 | Wild-type *B. subtilis* carrying the plasmid p8999-cas9-2N20-HA-169.5 | This study |
| p8999-cas9-bdhA-A3 | Wild-type *B. subtilis* carrying the plasmid p8999-cas9-bdhA-A3 | This study |
| p8999-cas9-bdhA-A4 | Wild-type *B. subtilis* carrying the plasmid p8999-cas9-bdhA-A4 | This study |
| p8999-cas9-bdhA-A7 | Wild-type *B. subtilis* carrying the plasmid p8999-cas9-bdhA-A7 | This study |
| p8999-cas9-bdhA-A15 | Wild-type *B. subtilis* carrying the plasmid p8999-cas9-bdhA-A15 | This study |
| p8999-cas9-bdhA-A37 | Wild-type *B. subtilis* carrying the plasmid p8999-cas9-bdhA-A37 | This study |
| p8999-cas9-bdhA-A38 | Wild-type *B. subtilis* carrying the plasmid p8999-cas9-bdhA-A38 | This study |
| p8999-cas9-bdhA-A44 | Wild-type *B. subtilis* carrying the plasmid p8999-cas9-bdhA-A44 | This study |
| p8999-cas9-bdhA-A49 | Wild-type *B. subtilis* carrying the plasmid p8999-cas9-bdhA-A49 | This study |
| pXY-dSpRY-A4 | Wild-type *B. subtilis* carrying the plasmid pXY-dSpRY-A4 | This study |
| pXY-dSpRY-A5 | Wild-type *B. subtilis* carrying the plasmid pXY-dSpRY-A5 | This study |
| pXY-dSpRY-A7 | Wild-type *B. subtilis* carrying the plasmid pXY-dSpRY-A7 | This study |
| pXY-dSpRY-A19 | Wild-type *B. subtilis* carrying the plasmid pXY-dSpRY-A19 | This study |
| pXY-dSpRY-A21 | Wild-type *B. subtilis* carrying the plasmid pXY-dSpRY-A21 | This study |
| pXY-dSpRY-A25 | Wild-type *B. subtilis* carrying the plasmid pXY-dSpRY-A25 | This study |
| pXY-dSpRY-A27 | Wild-type *B. subtilis* carrying the plasmid pXY-dSpRY-A27 | This study |
| pXY-dSpRY-A29 | Wild-type *B. subtilis* carrying the plasmid pXY-dSpRY-A29 | This study |
| pXY-dSpRY-A33 | Wild-type *B. subtilis* carrying the plasmid pXY-dSpRY-A33 | This study |
| pXY-dSpRY-A38 | Wild-type *B. subtilis* carrying the plasmid pXY-dSpRY-A38 | This study |
| pXY-dSpRY-A43 | Wild-type *B. subtilis* carrying the plasmid pXY-dSpRY-A43 | This study |
| pXY-dSpRY-A44 | Wild-type *B. subtilis* carrying the plasmid pXY-dSpRY-A44 | This study |
| pXY-dSpRY-A46 | Wild-type *B. subtilis* carrying the plasmid pXY-dSpRY-A46 | This study |
| pXY-dSpRY-A49 | Wild-type *B. subtilis* carrying the plasmid pXY-dSpRY-A49 | This study |
| p8999-cas9-nprE-wprA | Wild-type *B. subtilis* carrying the plasmid p8999-cas9-nprE-wprA | This study |
| p8999-cas9-wprA-epr | Wild-type *B. subtilis* carrying the plasmid p8999-cas9-wprA-epr | This study |
| p8999-cas9-17.8-19.6 | Wild-type *B. subtilis* carrying the plasmid p8999-cas9-17.8-19.6 | This study |
| p8999-cas9-19.6-23.3 | Wild-type *B. subtilis* carrying the plasmid p8999-cas9-19.6-23.3 | This study |
| p8999-cas9-wprA-epr-nprE | Wild-type *B. subtilis* carrying the plasmid p8999-cas9-wprA-epr-nprE | This study |
| p8999-cas9-SecY-A1 | Wild-type *B. subtilis* carrying the plasmid p8999-cas9-SecY-A1 | This study |
| p8999-cas9-SecY-A2 | Wild-type *B. subtilis* carrying the plasmid p8999-cas9-SecY-A2 | This study |
| p8999-cas9-SecY-A3 | Wild-type *B. subtilis* carrying the plasmid p8999-cas9-SecY-A3 | This study |
| p8999-cas9-SecY-A4 | Wild-type *B. subtilis* carrying the plasmid p8999-cas9-SecY-A4 | This study |
| p8999-cas9-SecY-A5 | Wild-type *B. subtilis* carrying the plasmid p8999-cas9-SecY-A5 | This study |
| p8999-cas9-SecY-A6 | Wild-type *B. subtilis* carrying the plasmid p8999-cas9-SecY-A6 | This study |
| p8999-cas9-SecY-A7 | Wild-type *B. subtilis* carrying the plasmid p8999-cas9-SecY-A7 | This study |
| p8999-cas9-SecY-A8 | Wild-type *B. subtilis* carrying the plasmid p8999-cas9-SecY-A8 | This study |
| p8999-cas9-SecY-A9 | Wild-type *B. subtilis* carrying the plasmid p8999-cas9-SecY-A9 | This study |
| p8999-cas9-SecY-A10 | Wild-type *B. subtilis* carrying the plasmid p8999-cas9-SecY-A10 | This study |
| p8999-cas9-SecY-A11 | Wild-type *B. subtilis* carrying the plasmid p8999-cas9-SecY-A11 | This study |
| p8999-cas9-SecY-A12 | Wild-type *B. subtilis* carrying the plasmid p8999-cas9-SecY-A12 | This study |
| p8999-cas9-SecY-A13 | Wild-type *B. subtilis* carrying the plasmid p8999-cas9-SecY-A13 | This study |
| p8999-cas9-SecY-A14 | Wild-type *B. subtilis* carrying the plasmid p8999-cas9-SecY-A14 | This study |
| p8999-cas9-SecY-A15 | Wild-type *B. subtilis* carrying the plasmid p8999-cas9-SecY-A15 | This study |
| p8999-cas9-SecY-A16 | Wild-type *B. subtilis* carrying the plasmid p8999-cas9-SecY-A16 | This study |
| p8999-cas9-SecY-A17 | Wild-type *B. subtilis* carrying the plasmid p8999-cas9-SecY-A17 | This study |
| p8999-cas9-SecY-A18 | Wild-type *B. subtilis* carrying the plasmid p8999-cas9-SecY-A18 | This study |
| p8999-cas9-SecY-A19 | Wild-type *B. subtilis* carrying the plasmid p8999-cas9-SecY-A19 | This study |
| p8999-cas9-SecY-A20 | Wild-type *B. subtilis* carrying the plasmid p8999-cas9-SecY-A20 | This study |
| p8999-cas9-SecY-A21 | Wild-type *B. subtilis* carrying the plasmid p8999-cas9-SecY-A21 | This study |
| p8999-cas9-SecY-A22 | Wild-type *B. subtilis* carrying the plasmid p8999-cas9-SecY-A22 | This study |
| p8999-cas9-SecY-A23 | Wild-type *B. subtilis* carrying the plasmid p8999-cas9-SecY-A23 | This study |
| p8999-cas9-SecY-A24 | Wild-type *B. subtilis* carrying the plasmid p8999-cas9-SecY-A24 | This study |
| p8999-cas9-SecY-A25 | Wild-type *B. subtilis* carrying the plasmid p8999-cas9-SecY-A25 | This study |
| p8999-cas9-SecY-A26 | Wild-type *B. subtilis* carrying the plasmid p8999-cas9-SecY-A26 | This study |
| p8999-cas9-SecY-A27 | Wild-type *B. subtilis* carrying the plasmid p8999-cas9-SecY-A27 | This study |
| p8999-cas9-SecY-A28 | Wild-type *B. subtilis* carrying the plasmid p8999-cas9-SecY-A28 | This study |
| p8999-cas9-SecY-A29 | Wild-type *B. subtilis* carrying the plasmid p8999-cas9-SecY-A29 | This study |
| p8999-cas9-SecY-A30 | Wild-type *B. subtilis* carrying the plasmid p8999-cas9-SecY-A30 | This study |
| p8999-cas9-SecY-A31 | Wild-type *B. subtilis* carrying the plasmid p8999-cas9-SecY-A31 | This study |
| p8999-cas9-SecY-A32 | Wild-type *B. subtilis* carrying the plasmid p8999-cas9-SecY-A32 | This study |
| p8999-cas9-SecY-A33 | Wild-type *B. subtilis* carrying the plasmid p8999-cas9-SecY-A33 | This study |
| p8999-cas9-SecY-A34 | Wild-type *B. subtilis* carrying the plasmid p8999-cas9-SecY-A34 | This study |
| p8999-cas9-SecY-A35 | Wild-type *B. subtilis* carrying the plasmid p8999-cas9-SecY-A35 | This study |
| p8999-cas9-SecY-A36 | Wild-type *B. subtilis* carrying the plasmid p8999-cas9-SecY-A36 | This study |
| p8999-cas9-SecY-A37 | Wild-type *B. subtilis* carrying the plasmid p8999-cas9-SecY-A37 | This study |
| p8999-cas9-SecY-A38 | Wild-type *B. subtilis* carrying the plasmid p8999-cas9-SecY-A38 | This study |
| p8999-cas9-SecY-A39 | Wild-type *B. subtilis* carrying the plasmid p8999-cas9-SecY-A39 | This study |
| p8999-cas9-SecY-A40 | Wild-type *B. subtilis* carrying the plasmid p8999-cas9-SecY-A40 | This study |
| p8999-cas9-SecY-A41 | Wild-type *B. subtilis* carrying the plasmid p8999-cas9-SecY-A41 | This study |
| p8999-cas9-SecY-A42 | Wild-type *B. subtilis* carrying the plasmid p8999-cas9-SecY-A42 | This study |
| p8999-cas9-SecY-A43 | Wild-type *B. subtilis* carrying the plasmid p8999-cas9-SecY-A43 | This study |
| p8999-cas9-SecY-A44 | Wild-type *B. subtilis* carrying the plasmid p8999-cas9-SecY-A44 | This study |
| p8999-cas9-SecY-A45 | Wild-type *B. subtilis* carrying the plasmid p8999-cas9-SecY-A45 | This study |
| p8999-cas9-SecY-A46 | Wild-type *B. subtilis* carrying the plasmid p8999-cas9-SecY-A46 | This study |
| p8999-cas9-SecY-A47 | Wild-type *B. subtilis* carrying the plasmid p8999-cas9-SecY-A47 | This study |
| p8999-cas9-SecY-A48 | Wild-type *B. subtilis* carrying the plasmid p8999-cas9-SecY-A48 | This study |
| p8999-cas9-SecY-A49 | Wild-type *B. subtilis* carrying the plasmid p8999-cas9-SecY-A49 | This study |
| p8999-cas9-SecY-A50 | Wild-type *B. subtilis* carrying the plasmid p8999-cas9-SecY-A50 | This study |

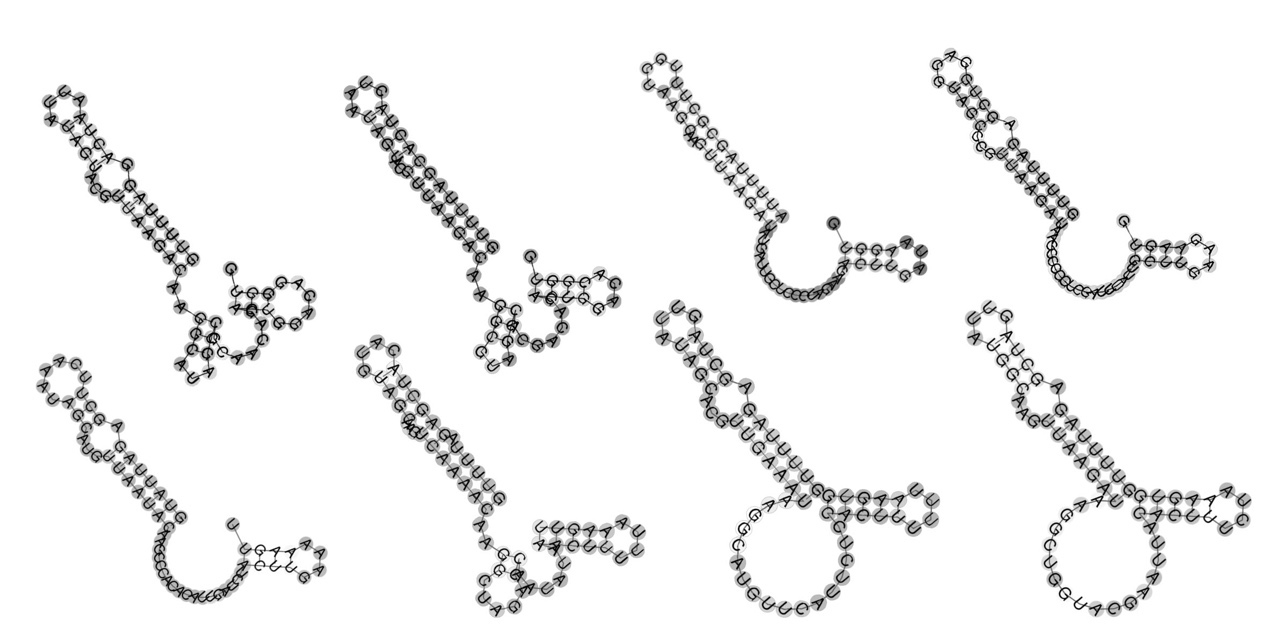

**Figure S1.** The secondary structure of the part of sgRNAs.

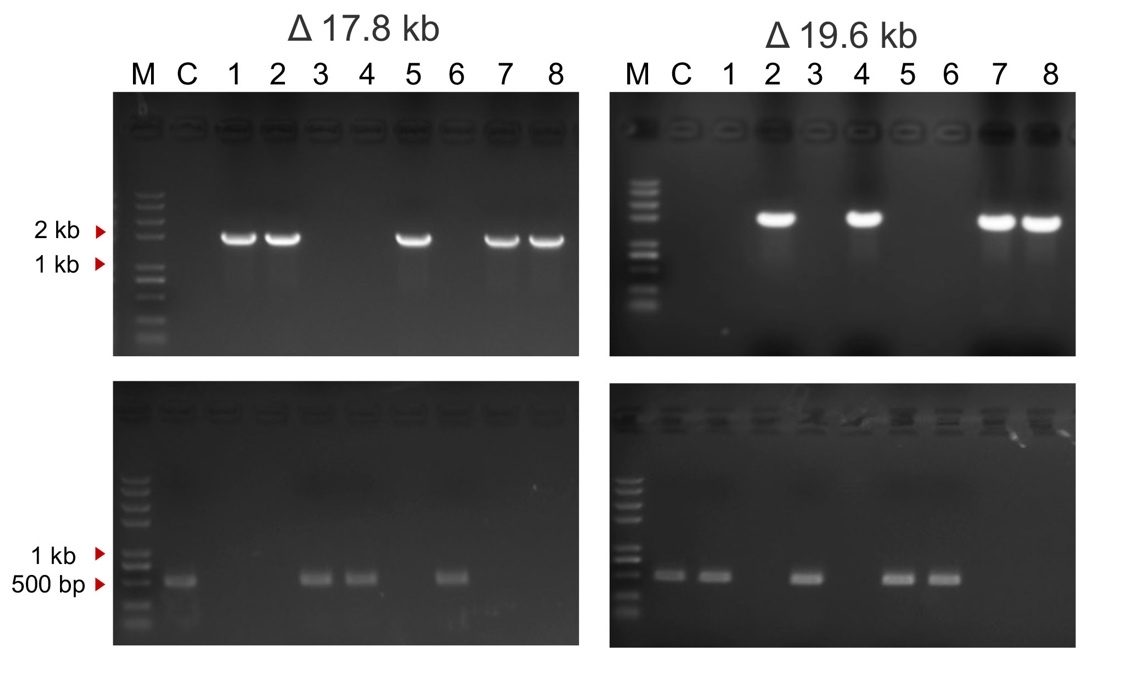

**Figure S2.** The editing results of large-scale knockout.

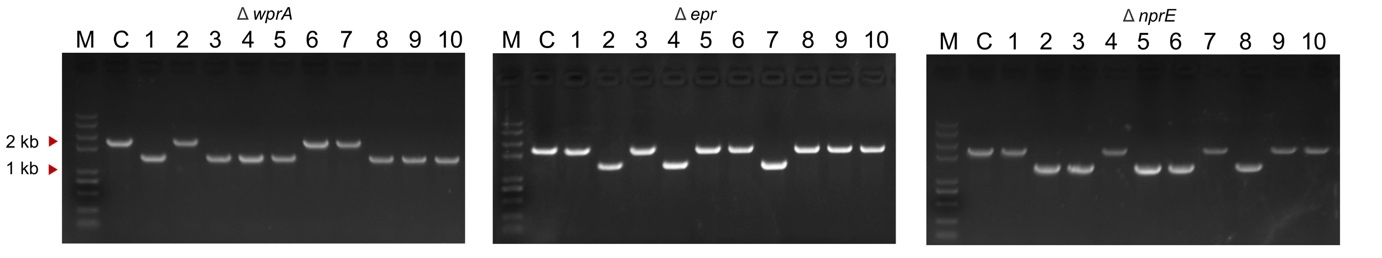

**Figure S3.** Editing results of simultaneous editing of three locis.

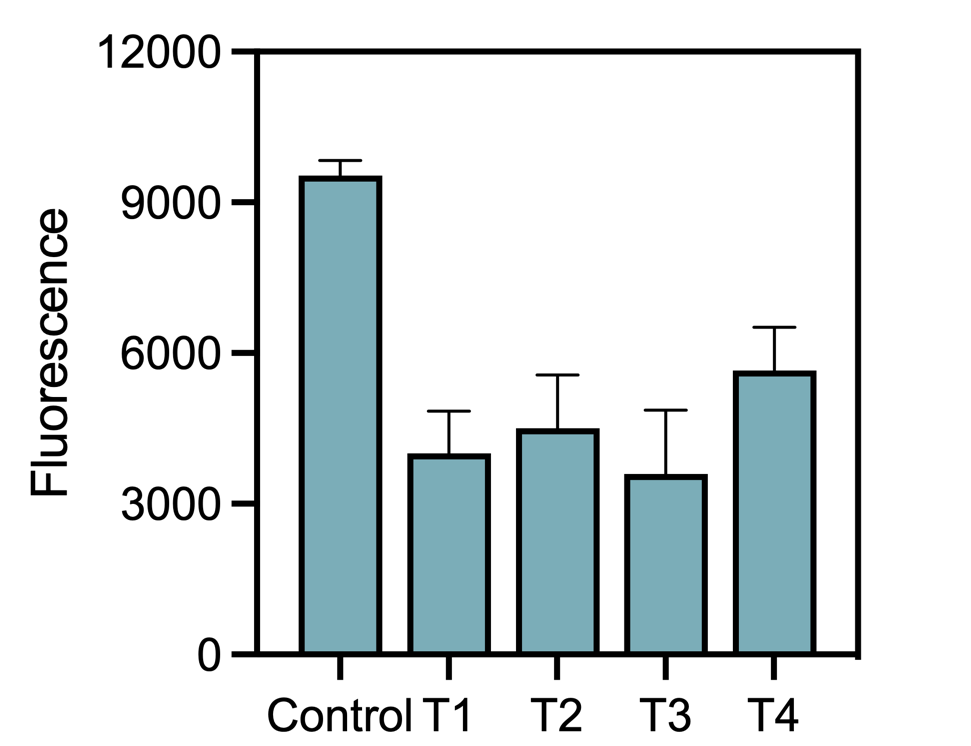

**Figure S4.** Fluorescence intensity across different target sites by repress different promoter region.
